# Wnt target IQGAP3 promotes Wnt signaling via disrupting Axin1-CK1α interaction

**DOI:** 10.1101/2024.12.10.626710

**Authors:** Muhammad Bakhait Rahmat, Aashiq Hussain, Teh Yu Xuan, Bibek Dutta, Sumedha Pundrik, Dennis Kappei, Yoshiaki Ito

## Abstract

The scaffold protein IQGAP3 is highly upregulated in most epithelial cancers. While recent studies have highlighted its pivotal roles in cancer cell proliferation and metastasis, a deeper mechanistic understanding of IQGAP3 is currently lacking. We have here used TurboID to map IQGAP3 proximity partners and identified the Wnt signaling members Axin1 and CK1α as IQGAP3-interacting proteins. Our functional studies demonstrated that overexpression of IQGAP3 increases β-catenin levels, while IQGAP3 depletion reduces β-catenin levels in gastric cancer cells. Mechanistically, IQGAP3 disrupts Axin1-CK1α interaction, thereby inhibiting β-catenin phosphorylation and ultimately leading to its accumulation. Importantly, we discovered that IQGAP3 itself is regulated by Wnt signaling, suggesting its involvement in a positive feedback loop in Wnt/β-catenin signaling through interactions with Axin1 and CK1α. These findings identify IQGAP3 as a novel mediator of β-catenin stabilization and underscore its potential as a target for cancer therapy.

## INTRODUCTION

IQ motif GTPase-activating scaffold proteins (IQGAPs) belong to a family of scaffold proteins, which facilitates physical interactions between effector proteins to coordinate signaling pathways involved in cytoskeleton dynamics (Brandt & Grosse, 2007, Wang, Watanabe et al., 2007), cell proliferation (Nojima, Adachi et al., 2008), and cytokinesis (Adachi, Kawasaki et al., 2014). The mammalian IQGAP family consists of three proteins: IQGAP1, IQGAP2, and IQGAP3. The prevalence of IQGAP1 and IQGAP3 in multiple cancer types indicates fundamental roles in tumorigenesis (Sanchez-Laorden, Viros et al., 2013), indeed, IQGAP1 and 3 have been established as potential targets for cancer therapy (Monteleon, McNeal et al., 2015).

IQGAP3 is located on chromosome 1q21.3, which is frequently amplified in many malignant tumors (Goh, Feng et al., 2017, Vazquez-Mena, Medina-Martinez et al., 2012). Amongst the IQGAP family, IQGAP3 alone is necessary and sufficient for driving cell proliferation (Nojima et al., 2008). Moreover, *Iqgap3* is specifically expressed in proliferating stomach stem cells and regenerating stomach tissues in the mouse (Matsuo, Douchi et al., 2021). The robust elevation of Iqgap3 following tissue injury is indicative of a requirement for IQGAP3 during wound healing (Matsuo et al., 2021). Injury-induced *Iqgap3* expression is associated upregulation of *Myc* gene targets, embryonic stem cell-like transcriptional signature and importantly, transcriptional programs associated with early gastric cancer (Matsuo, Douchi et al., 2020). More recently, the identification of oncofetal protein IGF2BP1 as a regulator of *IQGAP3* RNA stability suggests that fetal-like reprogramming in tumors led to IQGAP3 overexpression and hyperproliferation of stem-like cancer cells (Myint, Chuang et al., 2022). Mechanistically, IQGAP3 serves as an important hub for multiple oncogenic signaling pathways: PI3K/AKT (Lin, Liu et al., 2019), MAPK/ERK (Nojima et al., 2008, Yang, Zhao et al., 2014) and Wnt/β-catenin (Carmon, Gong et al., 2014). Understanding how IQGAP3 influences these pathways will help the design of effective strategies to inhibit proliferation and promote terminal differentiation in the ‘wound that does not heal’ – cancer.

Uncontrolled proliferation is a hallmark of cancer, driven by dysregulated signaling pathways that often involve scaffold and adaptor proteins to ensure precise spatiotemporal coordination. Recent findings highlight the role of biomolecular condensates formed via liquid-liquid phase separation (LLPS) as a key mechanism for coordinating intracellular processes (Mehta & Zhang, 2022, Nesterov, Ilyinsky et al., 2021, Tong, Tang et al., 2022), including cancer-related signaling (Hu, Wu et al., 2023, Meng, Yu et al., 2021). Proteins within LLPS function as scaffolds, driving phase separation, or clients, integrating into condensates to enhance multivalent interactions for efficient signal transduction (Banani, Rice et al., 2016). Wnt signaling, crucial for cell proliferation, differentiation, and stem cell maintenance, is frequently dysregulated in cancer, leading to β-catenin stabilization, overexpression of target genes (e.g., *MYC*, *CCND1*, *LGR5*) (Ramakrishnan & Cadigan, 2017), and subsequent tumor growth and metastasis. The β-catenin destruction complex, composed of Axin, APC, GSK3β, and CK1α, regulates β-catenin levels, with Axin’s phase separation playing a central role in complex formation and β-catenin phosphorylation. Disruption of Axin polymerization impairs LLPS formation (Bernkopf, Bruckner et al., 2019, Lach, Qiu et al., 2022), promoting Wnt signaling, while β-catenin itself undergoes phase separation via intrinsically disordered regions (IDRs), accumulating at super-enhancers to activate Wnt target genes (Zamudio, Dall’Agnese et al., 2019).

Our analysis of the IQGAP3 interactome provides an important resource for understanding the role of IQGAP3 in integrating signals from extensive protein networks in cancer. IQGAP3 functions as a scaffolding platform, supporting other scaffold proteins in coordinating complex signaling flux from multiple pathways. In this study, we identified Axin1 and CK1α as novel IQGAP3 interaction partner. We demonstrated that IQGAP3 promotes Wnt signaling by stabilizing β-catenin. Mechanistically, IQGAP3 reduces Axin1-CK1α interaction, thereby inhibiting β-catenin phosphorylation on Ser45 and leading to its accumulation. This global increase in β-catenin subsequently promotes the expression of pro-proliferation and survival genes. Finally, we identified an IQGAP3-Wnt feedback loop that sustains Wnt signaling.

## RESULTS

### Proximity proteomics reveal IQGAP3 interaction with Cdc42, Anillin and with centrosome- and septin-complex proteins

Since both IQGAP3 and Wnt signaling play crucial roles in the maintenance of gastric stem cells and carcinogenesis, we sought to understand the dynamics of IQGAP3 within the Wnt signal transduction network. Given the frequent overexpression of IQGAP3 in various cancer types, we utilized an IQGAP3 overexpression system to study its role in cancer progression. HEK293 cells were chosen for their intact Wnt signaling cascade (i.e., no mutations in Wnt signaling components) (Li, Ng et al., 2012), Wnt-responsiveness (Tuupanen, Turunen et al., 2009) and low basal expression of IQGAP3. We mapped the proximal interactome of IQGAP3 in living cells using the enzyme-catalyzed proximity labeling method, TurboID (Branon, Bosch et al., 2018) with and without Wnt3a stimulation. To induce Wnt signaling, HEK293 Tet-On cells were treated with L-Wnt3a cells (CRL-2647) conditioned media supplemented with rhWnt3a (100 ng/ml) or L-cells (CRL-2648) conditioned media (control) for 4 hours (Fig. 1A). To ensure that the TurboID tag did not compromise the expression, function, and localization of IQGAP3, we validated the construct in three ways: expression was confirmed through immunoblot analysis using both anti-HA-tag and anti-IQGAP3 antibodies (Fig. S1A), function was evaluated through anti-HA-tag immunoprecipitation and probing for Anillin, an established IQGAP3 interacting partner (Adachi et al., 2014) (Fig. S1B), and localization was assessed by immunofluorescence staining with anti-HA antibody (Fig. S1C).

**Fig. 1.**
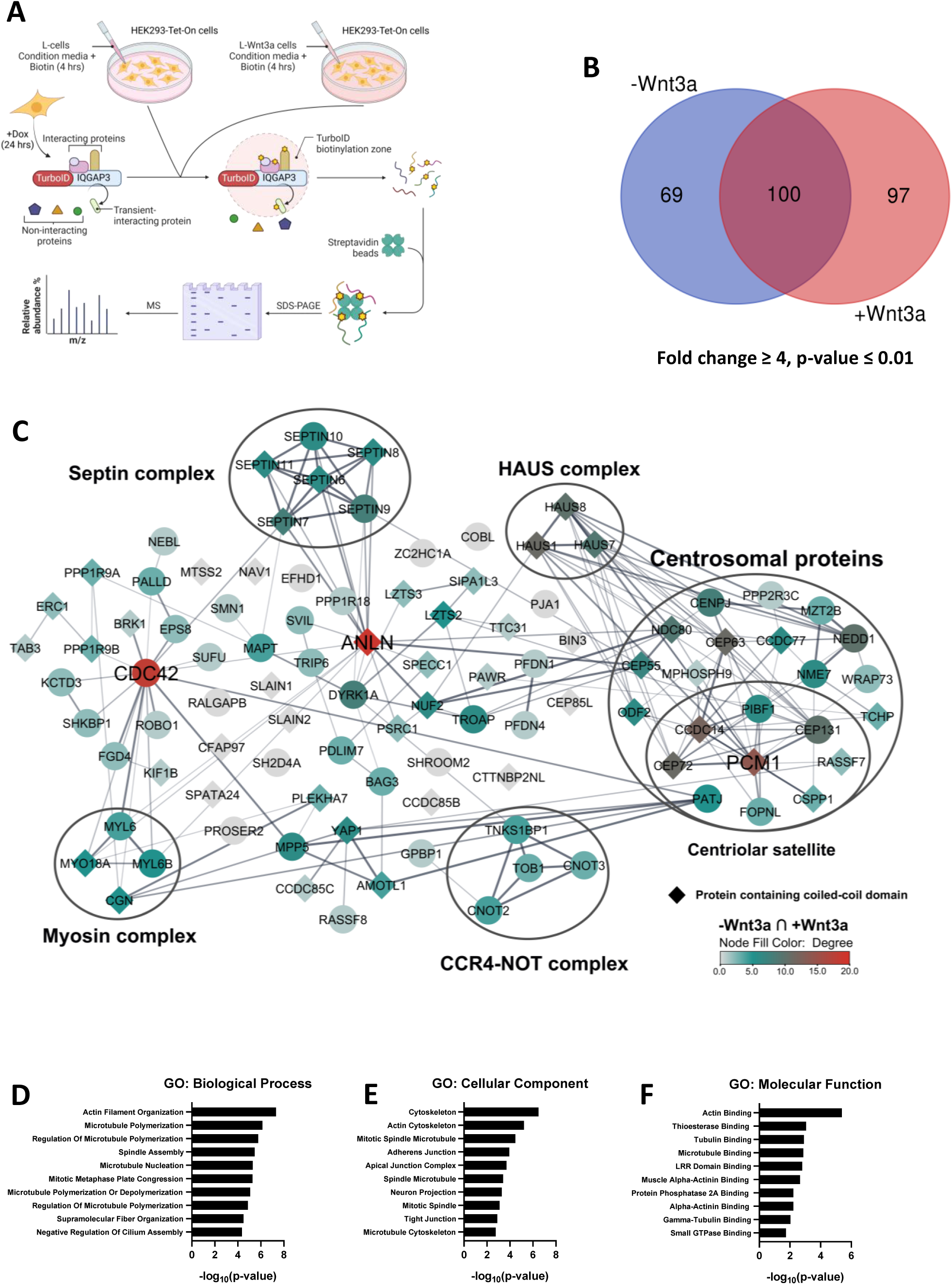
Proximity proteomics reveal IQGAP3 interaction with Cdc42, Anillin and with centrosome- and septin-complexes. (A) Workflow for proximity-dependent identification of IQGAP3-associated proteins in HEK293 cells treated with either L-cells condition media (-Wnt3a) or L-Wnt3a cells condition media (+Wnt3a) treatment for 4 hours, Created with BioRender.com. (B) Venn diagram of biotinylated proteins detected in HEK293 cells treated with L-cells condition media (-Wnt3a) and L-Wnt3a cells condition media (+Wnt3a) (Fold change ≥ 4, p-value ≤ 0.01). (C) Interactome of -Wnt3a ∩ +Wnt3a (Fold change ≥ 4, p-value ≤ 0.01), node color denote degree distribution, Created with Cytoscape. (D) Gene ontology: Biological Process of shared proximity partners in -Wnt3a and +Wnt3a treated cells. (E) Gene ontology: Molecular function of shared proximity partners in -Wnt3a and +Wnt3a treated cells. (F) Gene ontology: Cellular compartment of shared proximity partners in -Wnt3a and +Wnt3a treated cells.

Prior to analysis of TurboID-IQGAP3 by label-free quantitative mass spectrometry, protein expression and biotinylation levels were validated (Fig. S2A and B). 169 and 197 high-confidence IQGAP3 proximity partners were identified in the L-cells conditioned media (-Wnt3a) (Fig. S2C) and L-Wnt3a conditioned media (+Wnt3a) (Fig.S2D), respectively (Fold-change ≥ 4, p value ≤ 0.01), henceforth referred to as -Wnt3a and +Wnt3a treated cells. Amongst these, 100 proximity partners were found in both -Wnt3a and +Wnt3a treated cells (Fig 1B).

To systematically study the interactome profile changes brought about by Wnt3a treatment, we identified shared proximity partners present before and after Wnt3a treatment, exclusive proximity partners before and after Wnt3a treatment, and performed gene ontology analysis to further characterize their compartmentalization, biological and molecular functions.

Consistent with published literature, we observed established interacting partners of IQGAP3, such as Anillin (Adachi et al., 2014) and Cdc42 (Wang et al., 2007) in both interactomes (Fig. 1C). Interestingly, topological analysis of these established partners showed high degree (*k*) within the protein-protein interaction network, indicating their role as hub proteins i.e. proteins that are engaged in essential interactions (Barabasi, Gulbahce et al., 2011, Fox, Hescott et al., 2011, Stelzl, Worm et al., 2005). PCM1, a marker for centriolar satellites (Hori & Toda, 2017) was also identified as a strongly enriched proximity partner of IQGAP3 based on its high rank and degree within the interactomes (Fig. S1C and D).

Several complexes were also identified in both interactomes including Septin, CCR4-NOT, HAUS, and myosin complexes. Gene ontology for biological process, molecular function and cellular compartment indicates that the shared proximity partners of -Wnt3a and +Wnt3a treated cells are enriched for Actin Filament organization, Actin binding and Cytoskeleton, in congruence with its role as a regulator of actin cytoskeleton organization (Brandt & Grosse, 2007).

### Wnt3a treatment promotes IQGAP3 localization to non-membrane organelles and cell junctions

To gain insight into the changes to IQGAP3 proximity partners upon Wnt3a treatment, we compared the proximity partners that were exclusively enriched in the -Wnt3a and +Wnt3a treated cells. Exclusive proximity partners of IQGAP3 in -Wnt3a treated cells (n=69) revealed proteins involved in Wnt signaling; Axin1, APC, PPP2R1A and MACF1 (Fig 2A). Interestingly, all four proteins are scaffold proteins. The presence of IQGAP3 within the proximity of the destruction complex proteins, Axin1 and APC, indicates that it may play a role in Wnt-off condition. In addition, we again observed proteins involved in the CCR4-NOT and HAUS complex, which are known to be involved in mRNA metabolism (Denis & Chen, 2003) and centrosome/spindle integrity (Lawo, Bashkurov et al., 2009), respectively. Lastly, proximity partners found within P-bodies (i.e. dynamic non-membrane bound cytoplasmic structures) such as TOB1, EIF4ENIF1/4E-T, and TNRC6C, may suggest a role for IQGAP3 in the control of translation and mRNA degradation (Decker & Parker, 2012).

**Fig. 2.**
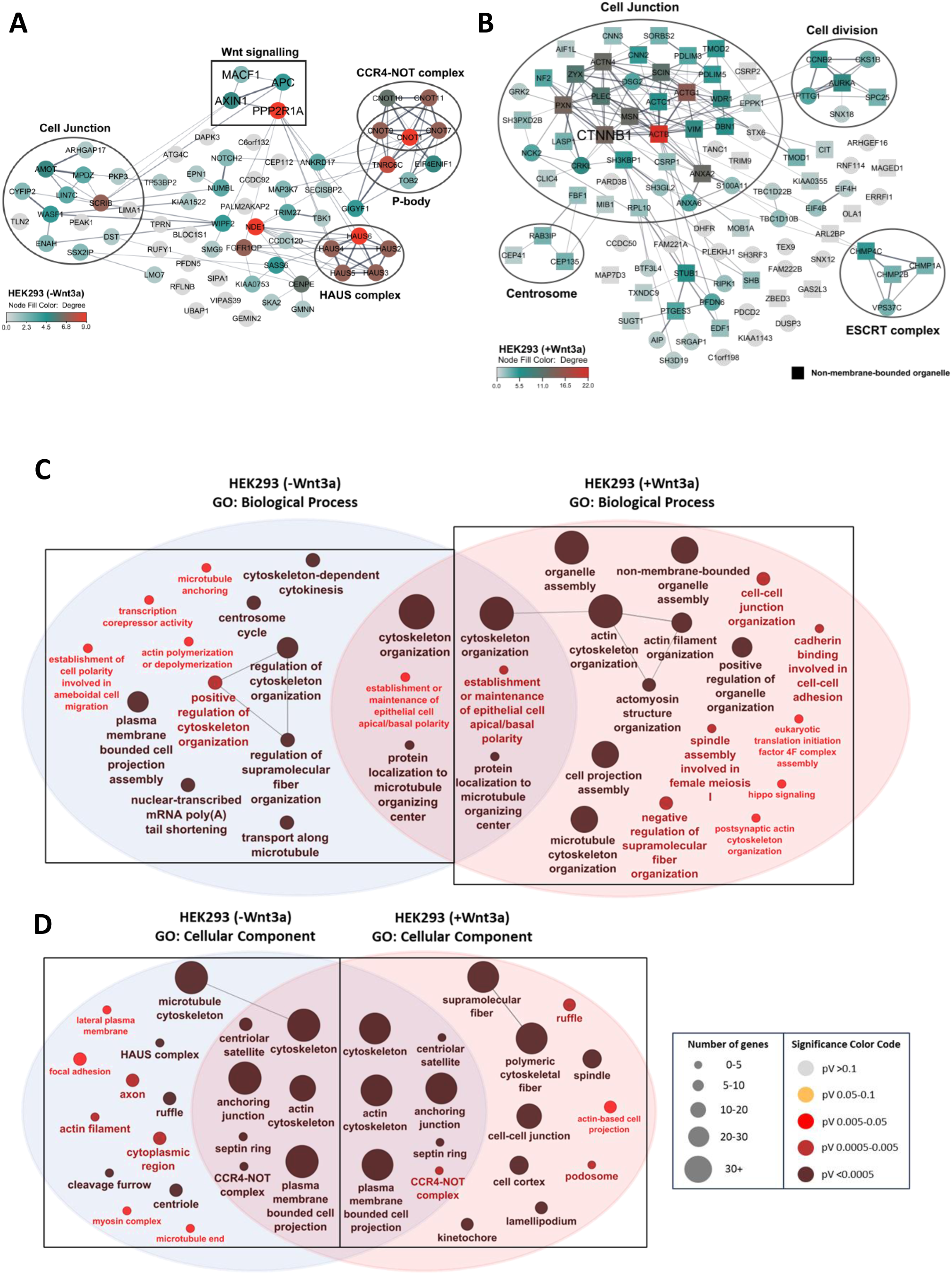
Wnt3a treatment promotes IQGAP3 localization to non-membrane organelles and cell junctions. (A) IQGAP3-biotinylated proteins exclusively found in cells treated with L-cells conditioned media (-Wnt3a). Shown are proximity partners with Fold change ≥ 4, p-value ≤ 0.01, node color denote degree distribution, Created with Cytoscape. (B) IQGAP3-biotinylated proteins exclusively found in cells treated with L-Wnt3a cells conditioned media (+Wnt3a). Shown are proximity partners with Fold change ≥ 4, p-value ≤ 0.01, node color denote degree distribution, Created with Cytoscape. (C) ClueGO analysis for Biological Process enrichment of biotinylated protein in HEK293 cells treated with -Wnt3a and +Wnt3a conditioned media. Size denote number of mapped genes and node color denote p-values. (D) ClueGO analysis for Cellular Compartment enrichment of biotinylated protein in HEK293 cells treated with -Wnt3a and +Wnt3a conditioned media. Size denote number of mapped genes and node color denote p-values.

Meanwhile, exclusive proximity partners of IQGAP3 in +Wnt3a treated cells (n=97) revealed increased in IQGAP3 proximity with cell junction proteins, ESCRT complex, proteins involved in cell division and the centrosome. Notably, we observed the loss of IQGAP3 proximity to Axin1, APC, and MACF1, along with an increased association with β-catenin and enrichment of proteins present in the cell junction, such as Paxillin, Zyxin, and Moesin (Fig. 2B).

Next, we compared the Gene ontology for Biological Process enriched in the -Wnt3a and +Wnt3a proximity partners. We observed that -Wnt3a treated cells showed enrichment for proximity partners involved in centrosome cycle, cell membrane projection, microtubule transport, cytoskeleton-dependent cytokinesis, regulation of cytoskeleton and supramolecular fiber organization. Meanwhile +Wnt3a treated cells showed enrichment for proximity partners involved in actin cytoskeleton organization, non-membrane bounded organelle-, organelle- and cell projection-assembly (Fig. 2C).

Comparison of the Gene ontology for Cellular Component revealed that -Wnt3a treated cells enriches proximity partners localized to the microtubule cytoskeleton, ruffle, cytoplasm, and centriole. Whereas +Wnt3a treated cells showed enrichment of proximity partners localized to the supramolecular fiber, cell-cell junction, lamellipodium, podosome and actin-based cell projection. Taken together, it appears that IQGAP3 responds to Wnt signaling by first associating with the destruction complex under Wnt-off conditions, and later translocating to the cell junction and actin-based cell projections upon stimulation with Wnt3a.

### IQGAP3 undergoes phase separation

To visualize changes in IQGAP3 localization within cells, we generated HeLa Tet-On cells stably expressing doxycycline-inducible EGFP-empty (control) or EGFP-IQGAP3 (Fig. 3A). HeLa cells were chosen because they are frequently used in imaging studies. Their large, flat morphology makes them ideal for high-resolution microscopy, as it facilitates the clear visualization of cellular structures, which is critical for protein localization studies.

**Fig. 3.**
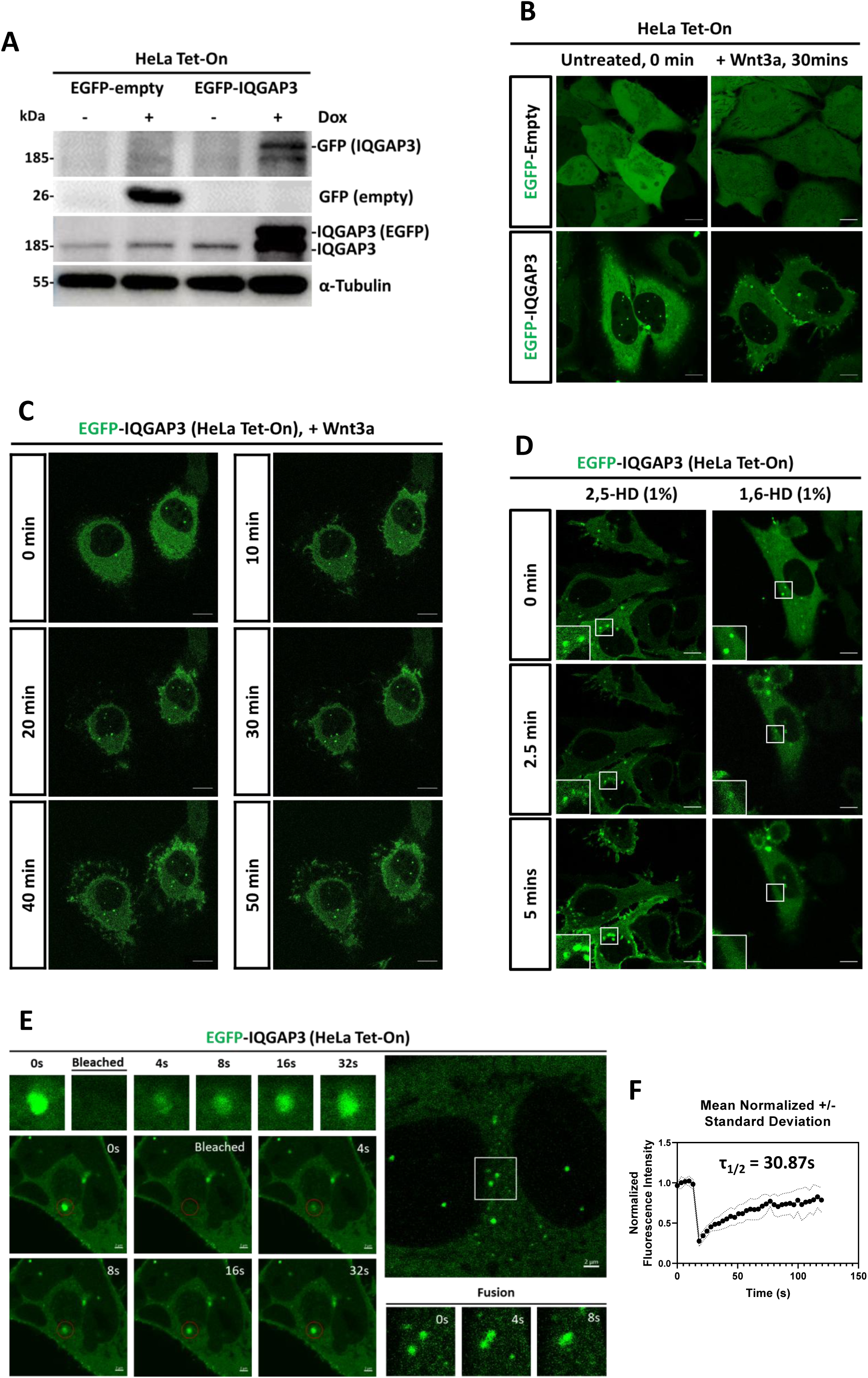
IQGAP3 undergoes phase separation. (A) Immunoblot analysis of Doxycycline induction in HeLa Tet-On cells. Cells were induced with Doxycycline for 48 hours prior to cell lysis. (B) Live images of HeLa Tet-On cells expressing EGFP-empty and EGFP-IQGAP3 without and with Wnt3a treatment after 30 minutes. Cells were induced with Doxycycline for 48 hours prior to imaging. Images of untreated and post-treated cells are not the same cells but are the best representative examples. Scale bars, 10 µm. (C) Timelapse images of HeLa Tet-On cells expressing EGFP-IQGAP3 0 - 50 minutes post-Wnt3a treatment. Cells were induced with Doxycycline for 48 hours prior to imaging. Images of untreated and post-treated cells are the same cells (refer to timelapse video in supplementary). Scale bars, 10 µm. (D) Timelapse images of HeLa Tet-On cells expressing EGFP-IQGAP3 treated with either 1% 2,5-Hexanediol (control) or 1% 1,6-Hexanediol for 0-5 minutes. Images of untreated and post-treated cells are the same cells (refer to timelapse video in supplementary). Scale bars, 10 µm. (E) HeLa Tet-On cells stably expressing EGFP-IQGAP3 were subjected to FRAP bleaching and observed for recovery and fusion events. (For FRAP video refer to supplementary). Scale bars, 2 µm. (F) Mean normalized standard deviation of FRAP recovery of 5 bleached EGFP-IQGAP3 condensates, half time of recovery (τ1/2) = 30.87s.

Timelapse experiments revealed the formation of IQGAP3 puncta in both the cytoplasm and nucleus (Fig. 3B and 3C). Furthermore, treatment of EGFP-IQGAP3 cells with Wnt3a resulted in an increased localization of IQGAP3 to the cell junctions, cell projections, and cytoplasmic puncta formation (Fig. 3C). Despite the differences in the cell lines used, these observations were consistent with the interactome data. Next, we treated the cells with 1% 1,6-hexanediol (1,6-HD), an aliphatic alcohol known to disrupt weak hydrophobic interactions and disassemble LLPS-dependent macromolecular condensates with minimal cytotoxicity (Alberti, Gladfelter et al., 2019). As a negative control, the cells were treated with 1% 2,5-Hexanediol (2,5-HD). To track the disruption of the puncta, we performed timelapse imaging before and after 1,6-HD and 2,5-HD treatments. We observed that while treatment with 1,6-HD disrupted EGFP-IQGAP3 condensates, treatment with 2,5-HD did not (Fig. 3D). We next examined whether IQGAP3 puncta display features distinctive of liquid-like condensates. A key feature of liquid-like condensates is their internal dynamical reorganization and fast exchange kinetics (Hyman, Weber et al., 2014), which can be assessed by measuring the rate of fluorescence recovery after photobleaching (FRAP). We studied the dynamics of EGFP-IQGAP3 condensates by FRAP and observed that EGFP-IQGAP3 displayed dynamic condensate formation distinctive of liquid-like condensates capable of both recovery after bleaching (half time of recovery, τ_1/2_ = 30.87s) and fusion with other condensates (Fig. 3E and F).

### IQGAP3 co-localizes within Axin1 puncta in the cytosol

Both IQGAP1 and IQGAP3 have been implicated to be involved in Wnt signaling (Carmon et al., 2014). To compare the localization of IQGAP1 and IQGAP3, we overexpressed full length IQGAP1 and IQGAP3 fused to mCherry2 in HeLa cells (Fig. 4B and 4C), and as control mcherry2-empty (Fig. 4A). We counted 50 cells that showed mCherry2-IQGAP1 or mCherry2-IQGAP3 expression and categorized them based on the distribution of the IQGAP proteins i.e. when the highest intensity of IQGAP expression is in the membrane, the cell is counted as Mem>Cyto, vice versa (Fig. 4D). We observed that IQGAP1 is predominantly localized to the membrane (Fig. 4B) whereas IQGAP3 is localized both in the cytoplasm and nucleus (Fig 4C).

**Fig. 4.**
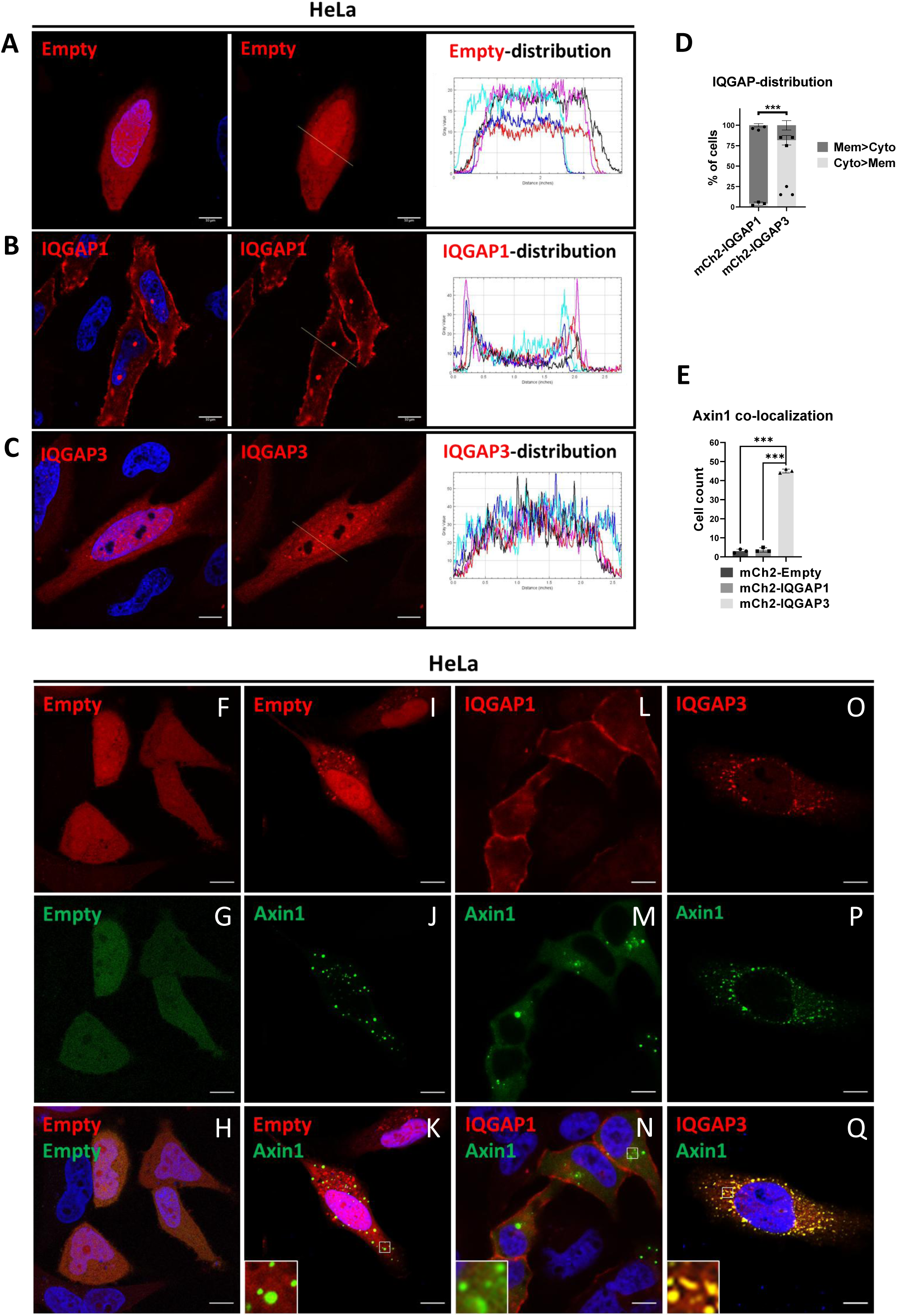
IQGAP3 co-localizes with Axin1 puncta in the cytosol. HeLa cells were transfected with (A) mCherry2-empty and Cross-sectional line for Intensity plot measurement from 5 different cells (right). (B) mCherry-IQGAP1 and Cross-sectional line for Intensity plot measurement from 5 different cells (right). (C) mCherry2-IQGAP3 and Cross-sectional line for Intensity plot measurement from 5 different cells (right). Scale bars, 10 µm. (D) Percentile of cell count for IQGAP-distribution, cells with higher membrane intensity were counted as Membrane > Cytosol (Mem > Cyto) vice versa. Cell count for IQGAP distribution, 50 cells per sample. Representative data were collected and are expressed as the mean ± SD from three independent experiments (n=3). (E) Cell count for IQGAP-Axin1 co-localization, 50 cells per sample. Representative data were collected and are expressed as the mean ± SD from three independent experiments (n=3). Student’s t test was performed, with ∗p < 0.05, ∗∗p < 0.01, and ∗∗∗p < 0.001. HeLa cells were transfected with (F-H) mCherry2-empty with EGFP-empty (negative control), (I-K) mCherry2-empty with EGFP-Axin1, (L-N) mCherry2-IQGAP1 with EGFP-Axin1, and (O-Q) mCherry2-IQGAP3 with EGFP-Axin1. Scale bars, 10 µm.

Axin1 is a proximity partner of IQGAP3 in the HEK293 interactome. Importantly, Axin1 is known to form LLPS and its puncta formation is correlated with the activity of the β-catenin destruction complex (Bernkopf et al., 2019, Nong, Kang et al., 2021, Schaefer & Peifer, 2019, Shi, Kang et al., 2021). To study this phenomenon, we employed the HeLa cell line which is commonly used to model biomolecular condensate formation *in vivo* (Muzzopappa, Hummert et al., 2022, Poudyal, Patel et al., 2023, Shakya, Park et al., 2020). As controls, we expressed mCherry2-empty with EGFP-empty (Fig. 4F-H) and mCherry2-empty with EGFP-Axin1 (Fig. 4I-K). Ectopic expression of EGFP-Axin1 with mCherry2-IQGAP1 (Fig. 4L-N) or mCherry2-IQGAP3 (Fig. 4O-Q) revealed only the colocalization of IQGAP3 within Axin1 puncta. Furthermore, exogenous co-immunoprecipitation of IQGAP1 and IQGAP3 revealed that only IQGAP3 binds Axin1 (Fig. S3A and B). Taken together, the differences in their subcellular distribution may explain the differential binding of IQGAP1 and IQGAP3 to Axin1. Therefore, although both paralogs have been linked to Wnt signaling (Carmon et al., 2014), they likely contribute to distinct aspects of the Wnt signaling cascade.

### IQGAP3 expression reduces Axin1-CK1α interaction

Next, we confirm the endogenous interaction between IQGAP3 and the destruction complex. Endogenous immunoprecipitation of IQGAP3 from HEK293 lysate revealed endogenous IQGAP3 interaction with Axin1, β-catenin, CK1α and PCM1 (Fig. 5A). IQGAP3 interaction with Axin1, β-catenin, and CK1α confirms its presence within the destruction complex. IQGAP3 however did not directly interact with other components of the destruction complex; GSK3β and PP2Aα. Similarly, exogenous co-immunoprecipitation experiments involving IQGAP3 with either Axin1, CK1α, GSK3β, PP2Ac, or β-catenin revealed that only Axin1 and CK1α binds to IQGAP3 (Fig. S3B-F), thus indicating the specificity of the interaction.

**Fig. 5.**
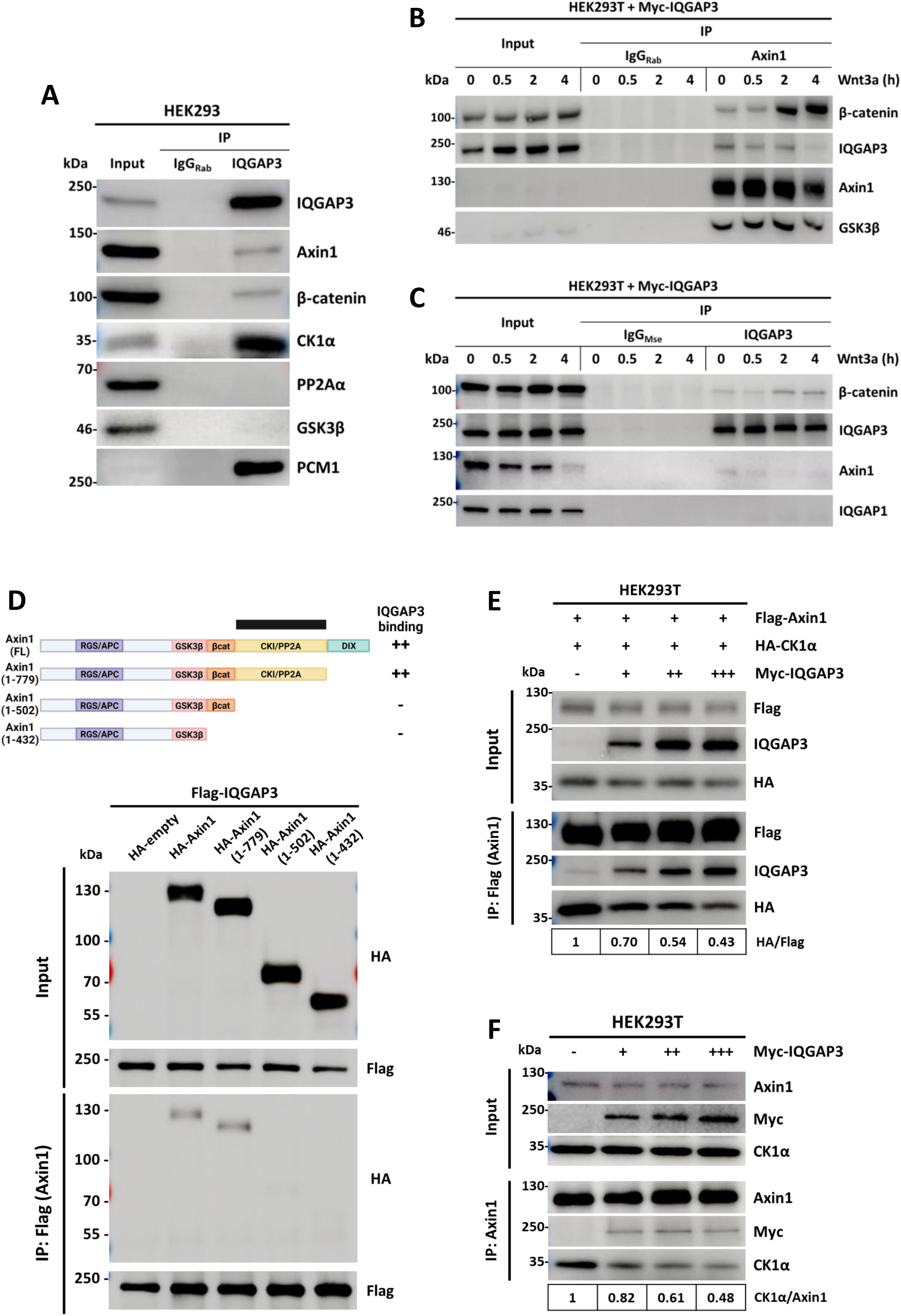
IQGAP3 expression reduces Axin1-CK1α interaction. (A) Immunoblot of endogenous IQGAP3 immunoprecipitation using an IQGAP3-specific antibody (1:100) and Mouse IgG as a control, with 1 mg of HEK293 cell lysate. Samples were subjected to SDS-PAGE (7.5%) followed by immunoblotting using the indicated antibodies. (B) HEK293T cells were exposed to Wnt3a conditioned medium at different time points as indicated. Stimulated lysates were subjected to immunoprecipitation using an Axin1-specific antibody and Rabbit IgG as control and were subjected to SDS-PAGE (7.5%) followed by immunoblotting using the indicated antibodies. (C) HEK293T cells were exposed to Wnt3a conditioned medium at different time points as indicated. Stimulated lysates were subjected to IQGAP3 immunoprecipitation using Myc-Tag antibody and Mouse IgG as control and were subjected to SDS-PAGE (7.5%) followed by immunoblotting using the indicated antibodies. (D) Immunoblot of Flag-Immunoprecipitated protein expressing HA-Axin1 C-terminal truncations with Flag-IQGAP3 to map the IQGAP3 binding site within Axin1. Samples were subjected to SDS-PAGE (7.5%) followed by immunoblotting using the indicated antibodies. (E) IQGAP3 reduced exogenous Axin1-CK1 interaction. HEK293T cells expressing Flag-Axin1 were transfected with limiting amounts of HA-CK1α and increasing amounts of Myc-IQGAP3 (5µg/10µg/15µg) as indicated, and the lysates were subjected to anti-Flag immunoprecipitation. Samples were subjected to SDS-PAGE (7.5%) followed by immunoblotting using the indicated antibodies. The relative immunoblot bands of HA and Flag (HA/Flag) from the immunoprecipitation membrane were quantified by densitometry. (F) IQGAP3 reduced endogenous Axin1-CK1 interaction. HEK293T cells were transfected with increasing amounts of Myc-IQGAP3 (5µg/10µg/15µg) as indicated, and the lysates were subjected to anti-Axin1 immunoprecipitation. Samples were subjected to SDS-PAGE (10%) followed by immunoblotting using the indicated antibodies. The relative immunoblot bands of CK1α and Axin1 (CK1α/Axin1) from the immunoprecipitation membrane were quantified by densitometry.

To study the dynamics of IQGAP3-Axin1 interaction, we performed a time-course Wnt3a treatment and performed immunoprecipitation using Axin1 antibody in HEK293T cells overexpressing Myc-IQGAP3 (Fig. 5B). We note that IQGAP3-Axin1 complex formation decreases at 4 hours post Wnt3a treatment due to less Axin1 precipitating as a result of Axin1 degradation that occurs at 4 hour post Wnt3a treatment as others have found (Li et al., 2012). This observation explained the loss of Axin1 from the interactome following Wnt3a treatment (Fig. 2B).

As expected, Wnt3a treatment showed increased Axin1-β-catenin complex formation with the duration of stimulation (Li et al., 2012). Similarly, immunoprecipitation using IQGAP3 antibody (Fig. 5C) showed increased IQGAP3-β-catenin complex formation with the duration of Wnt3a treatment consistent with the observation that β-catenin interaction is gained in the IQGAP3 interactome following Wnt3a treatment (Fig. 2B).

To characterize the Axin1-IQGAP3 interaction, we mapped the binding domains and found that IQGAP3 binds to the region between the β-catenin binding domain and the DIX domain (i.e., the CK1α/PP2A binding site) of Axin1 (Fig. 5D) via its IQ and RGCT domains (Fig. S3G-I). Given that IQGAP3 can bind to both CK1α and the CK1α/PP2A binding site within Axin1, we hypothesized that it might disrupt the Axin1-CK1α interaction. Indeed, increased expression of IQGAP3 in HEK293T cells reduced the presence of CK1α in both exogenous Axin1 (Fig. 5E) and endogenous Axin1 (Fig. 5F) precipitates, confirming that IQGAP3 overexpression reduces the Axin1-CK1α interaction. Although we observed that IQGAP3 interacts with Axin1, potentially blocking the Axin1-CK1α interaction, we cannot rule out the possibility that overexpressed IQGAP3 also sequesters CK1α away from Axin1, thereby simultaneously reducing the Axin1-CK1α interaction.

### IQGAP3 promotes β-catenin activity through its CC and IQ domains

We next examined the effect of IQGAP3 overexpression on Wnt signaling. Considering that β-catenin is necessary for Wnt activation and that Axin1 is a critical component of β-catenin destruction complex, we hypothesized that IQGAP3 interferes with β-catenin destruction by reducing the Axin1-CK1α interaction. To test this, we assessed the effects of IQGAP3 depletion and overexpression on exogenous β-catenin in HEK293T cells. Since free excess β-catenin is rapidly phosphorylated by the destruction complex and marked for degradation (Valenta, Hausmann et al., 2012), we used exogenous β-catenin expression to determine whether changes in IQGAP3 could affect this process (Goentoro & Kirschner, 2009).

To measure β-catenin activity we perform TOPFlash assay by co-expressing β-catenin with either Axin1 or an increasing amount of IQGAP3. We observed that β-catenin co-expression with Axin1 drastically reduced β-catenin activity, while co-expression with increasing amount of IQGAP3 led to a significant increase in β-catenin activity (Fig. 6A). In contrast, β-catenin co-expression with increasing amount of siIQGAP3 reduced β-catenin activity (Fig. 6B). Validation for siIQGAP3 knockdown (Fig. S4H).

**Fig. 6.**
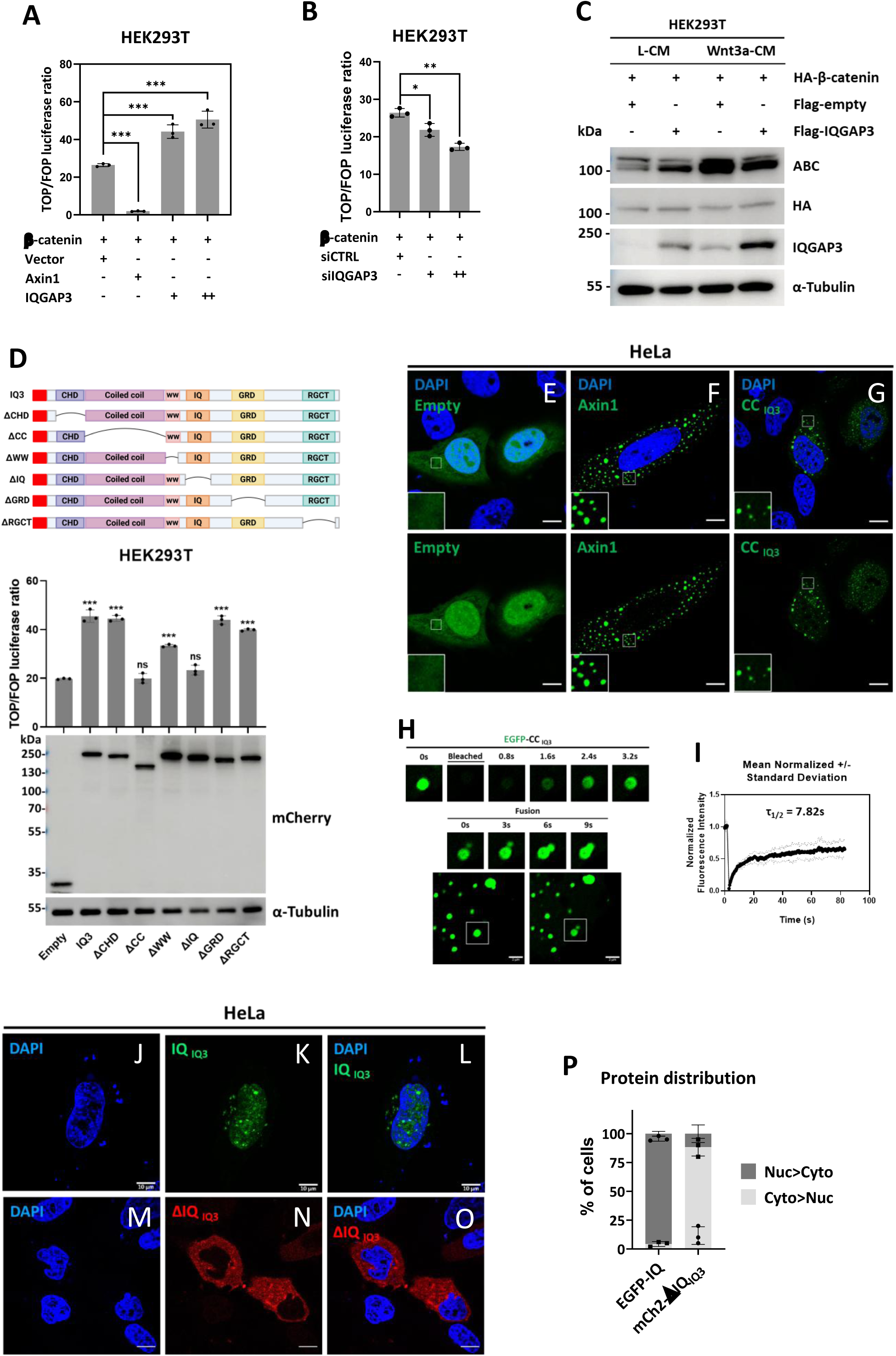
IQGAP3 promotes β-catenin activity through its CC and IQ domains. (A) TOPFlash assay of β-catenin co-overexpression with Axin1 or increasing IQGAP3 (300µg, 600µg) in HEK293T cells. The data is representative of three independent experiments. Student’s t test was performed, with ∗p < 0.05, ∗∗p < 0.01, and ∗∗∗p < 0.001. (B) TOPFlash assay of β-catenin co-overexpression with siControl or increasing siIQGAP3 (20nM, 50nM) in HEK293T cells. The data is representative of three independent experiments. Student’s t test was performed, with ∗p < 0.05, ∗∗p < 0.01, and ∗∗∗p < 0.001. (C) Immunoblot of IQGAP3 overexpression with and without Wnt3a conditioned media treatment. The data is representative of three independent experiments. (D) TOPFlash and Immunoblot assays of β-catenin overexpression with a series of IQGAP3 domain deletions. The data is representative of three independent experiments. HeLa cells transfected with (E) EGFP-empty, (F) EGFP-Axin1, and (G) EGFP-CC. Scale bars, 10 µm. (H) HeLa cells were transfected with EGFP-CC, subjected to FRAP bleaching, and observed for recovery and fusion events. Scale bars, 2µm. (I) Mean normalized standard deviation of FRAP recovery of 10 bleached EGFP-CC condensates, half time of recovery (τ1/2) = 7.82s. HeLa cells transfected with (J-L) EGFP-IQ domain and (M-O) mCherry2-IQGAP3ΔIQ. Scale bars, 10 µm. (P) Percentile of cell count for protein distribution, cells with higher nuclear intensity were counted as Nuclear > Cytosol (Nuc > Cyto), vice versa. Cell count for protein distribution, 50 cells per sample. Representative data were collected and are expressed as the mean ± SD from three independent experiments (n=3).

The priming phosphorylation on Serine 45 of β-catenin is mediated by CK1α and facilitated by Axin1. (Marin, Bustos et al., 2003). Therefore, disruption of Axin1-CK1α interaction will affect S45 phosphorylation and eventually β-catenin levels. Consistent with this, exogenous expressions of IQGAP3 increased non-phosphorylated β-catenin (at Ser45) (hereafter, referred to as ABC), in Wnt-off condition but decreased ABC level in Wnt-on condition (Fig. 6C). Hence, overexpressed IQGAP3 may positively regulate β-catenin activity under Wnt-off conditions; however, it may negatively regulate β-catenin activity under Wnt-on conditions. This dual role of IQGAP3 in Wnt signaling may be explained by the interactome data, which shows that Wnt3a treatment, combined with IQGAP3 overexpression, causes IQGAP3 to relocate to the cell junction with β-catenin. Interestingly, we also observed that Wnt3a treatment increases IQGAP3 expression.

To identify the IQGAP3 domain involved in promoting β-catenin activity, we perform TOPFlash assay with a series of IQGAP3 domain deletion constructs (Fig. 6D). Deletion of the Calponin homology domain (ΔCHD), WW domain (ΔWW), GAP-related domain (ΔGRD), and RasGAP C-terminal (ΔRGCT) promoted β-catenin activity (Fig. 6C lanes 3, 5, 7 and 8), whereas deletion of the Coiled coil (ΔCC) and IQ domain (ΔIQ) failed to promote β-catenin activity (Fig. 6D lanes 4 and 6). We concluded that both CC and IQ domains are required for promoting β-catenin activity.

To gain insight into the function of these domains we amplified the cDNA encoding for the coiled coil (L147 and F669) and IQ (V734 to P849) domains and cloned them into the EGFP-empty vector. We then overexpressed EGFP-coiled coil (CC) and EGFP-IQ in HeLa cells. We observed that EGFP-CC generated distinct puncta formation (Fig. 6G). Similarly, when we studied the dynamics of EGFP-CC condensates using FRAP, we found that EGFP-CC exhibited even higher dynamics in condensate formation after bleaching with a half time of recovery at τ_1/2_=7.42s compared to EGFP-IQGAP3 at τ_1/2_=30.87s (Fig. 6G-I). In addition, IQGAP3 deletion of various domains (Fig. S4A-G) showed loss of puncta formation when cell was transfected with the mCherry2-IQGAP3ΔCC construct (Fig. S4C).

Meanwhile, cells overexpressed with EGFP-IQ displayed mainly nuclear localization of the peptide (Fig. 6I-K). To confirm that IQ domain is sufficient and necessary for nuclear localization we overexpressed IQGAP3 with an IQ domain deletion (mCherry2-IQGAP3ΔIQ). We observed that IQ deletion abolished any presence of nuclear IQGAP3 (Fig.6L-N) compared to the deletions of other domains (Fig.S4A-G), therefore, the IQ domain within IQGAP3 may harbor a nuclear localization signal (NLS) required for its nuclear localization. To predict the presence of an NLS, we performed an *in silico* analysis of the IQGAP3 protein sequence using NLStradamus (Nguyen Ba, Pogoutse et al., 2009) and cNLS Mapper (Kosugi, Hasebe et al., 2009). Neither analysis predicted an NLS within IQGAP3. Therefore, we hypothesized that the IQ domain might instead bind to a protein with an NLS, mediating IQGAP3’s nuclear translocation—essentially ‘hitchhiking’ into the nucleus. Taken together, the CC and IQ domains mediate IQGAP3 phase separation and nuclear localization, respectively. Therefore, it appears that the ability of IQGAP3 to promote β-catenin activity is dependent on its ability to phase separate and translocate to the nucleus.

### IQGAP3 overexpression increases β-catenin levels in MKN28 cells

To determine IQGAP3-low and-high expressing gastric cancer cell lines, we screened several gastric cancer cell lines for the expression of IQGAP3 (Fig. 7A). NUGC3 and AGS cells showed high IQGAP3 expression, whereas MKN28 cells showed low IQGAP3 expression, hence, the latter was deemed a suitable candidate to study the effect of IQGAP3 overexpression in gastric cancer cells. We generated MKN28 cells stably overexpressing doxycycline-inducible 3×Flag-IQGAP3 and upon treatment with doxycycline showed increased β-catenin levels (Fig. 7B).

**Fig. 7.**
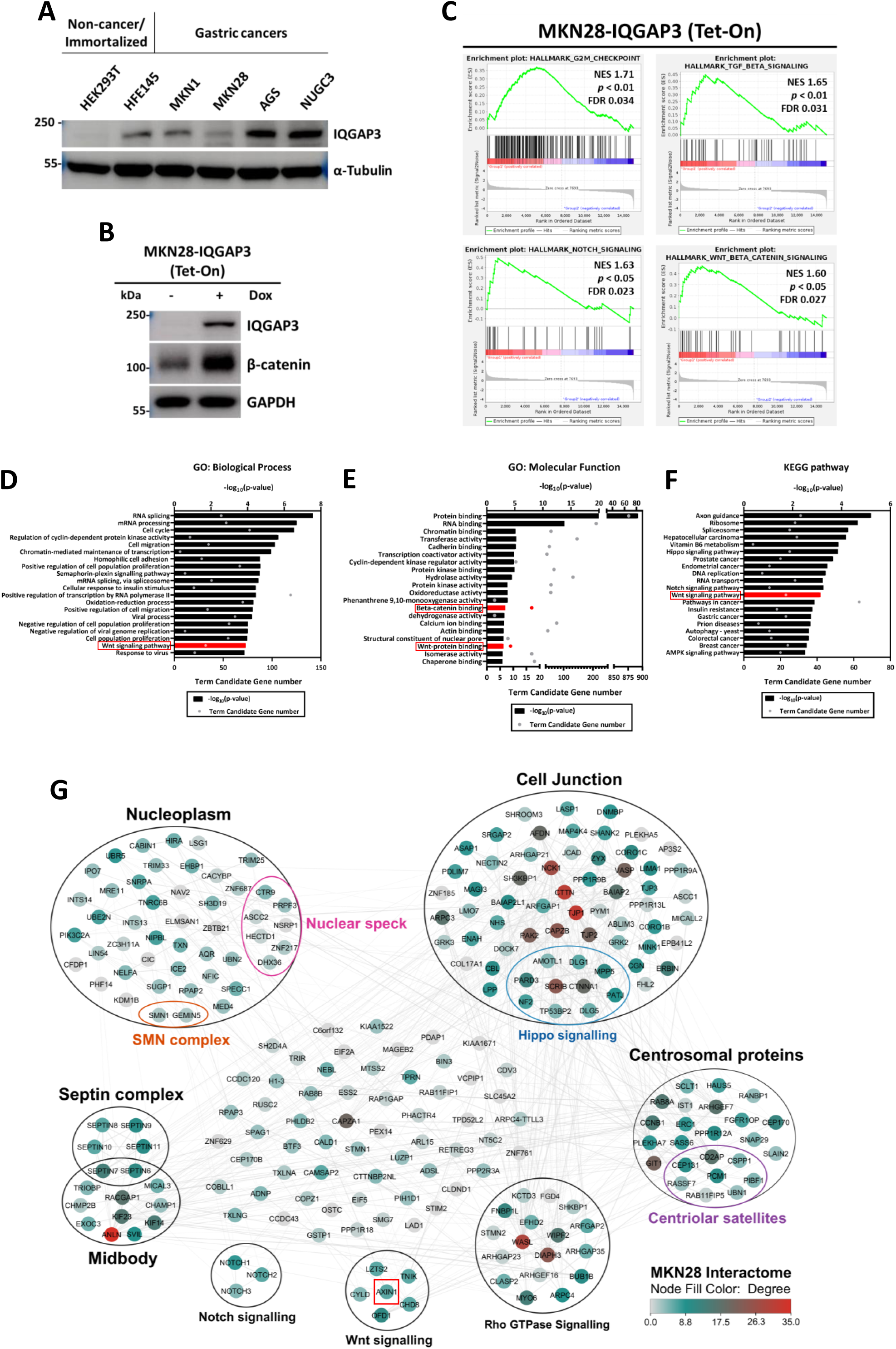
IQGAP3 overexpression increases β-catenin levels in MKN28 cells. (A) Immunoblot of non-cancer/immortalized (HEK293T and HFE145) and gastric cancer cell lines (MKN1, AGS, and NUGC3). Samples were subjected to SDS-PAGE (7.5%) followed by immunoblotting using the indicated antibodies. (B) Immunoblot of MKN28 cells expressing doxycycline-inducible 3×Flag-IQGAP3. Samples were subjected to SDS-PAGE (7.5%) followed by immunoblotting using the indicated antibodies. (C) GSEA data from RNA-seq of IQGAP3 overexpression in MKN28 cells, showing top 4 pathways. (D) Gene ontology: Biological Process Enrichment histogram of RNA-seq data from IQGAP3 overexpression in MKN28 cells. Wnt Signaling pathway represented with red bar. (E) Gene ontology: Molecular function Enrichment histogram of RNA-seq data from IQGAP3 overexpression in MKN28 cells. Wnt-protein and beta-catenin binding are represented with red bars. (F) KEGG pathway Enrichment histogram of RNA-seq data from IQGAP3 overexpression in MKN28 cells. Wnt Signaling pathway represented with red bar. (G) IQGAP3 interactome mapped from MKN28 cells stably expressing doxycycline-inducible TurboID-IQGAP3 (Fold change ≥ 2, p-value ≤ 0.05), node color denote degree distribution, Created with Cytoscape.

Next, we generated MKN28 cells stably overexpressing doxycycline-inducible 3×HA-TurboID-IQGAP3 for RNA-seq analysis and proximity ligation. Induction of fusion protein expressions and biotinylating activities were analyzed by immunoblot (Fig. S5A and S5B). The RNA-seq analysis from IQGAP3 overexpressing cells revealed an enrichment for several signaling pathways, G2/M, TGF-β, NOTCH, and Wnt/β-catenin in the GSEA data (Fig. 7C). Similarly, the gene ontology analysis of the Biological Process (GO) revealed enrichment in the Wnt signaling pathway (Fig. 7D), the Molecular Function (GO) showed enrichment in Wnt-protein and beta-catenin binding (Fig. 7E), and the KEGG pathway analysis indicated enrichment in Hippo, Notch and Wnt signaling pathways (Fig. 7F). For the full list of GSEA rankings from RNA-sequencing analysis see Fig. S5C. We also validated the MKN28 cells for IQGAP3-Axin1 interaction and changes in Wnt target genes (*Axin2* and *TCF7*) expression upon IQGAP3 overexpression via immunoprecipitation and RT-qPCR, respectively (Fig. S5E and F).

Next, we performed the biotinylation ligation of IQGAP3 proximity partners in MKN28 cells. Prior to analysis of TurboID-IQGAP3 by label-free quantitative mass spectrometry, protein expression and biotinylation levels were validated (Fig. S5A and B). 232 high-confidence IQGAP3 proximity partners were identified (Fold-change ≥ 2, p value ≤ 0.05) (Fig. 7G). Proximity ligation of IQGAP3 within MKN28 cells revealed Axin1 as a proximity partner of IQGAP3. The presence of IQGAP3 in proximity to Axin1 in both HEK293 and MKN28 cells strongly suggests its involvement in the destruction complex. Moreover, proteins involved in cell junction, septin complex, and centrosome are also found in the MKN28 interactome.

Surprisingly, we observed a greater number of nuclear proteins (nucleoplasm proteins) as IQGAP3 proximity partners in MKN28 cells compared to HEK293 cells. Among these nuclear proteins, seven (CTR9, PRPF3, ASCC2, NSRP1, HECTD1, ZNF217, and DHX36) were associated with nuclear speckles formation. Similarly, when analyzing endogenous IQGAP3 localization in non-transformed cell lines (HEK293T and HFE145) and transformed gastric cell lines (MKN1, MKN28, HS746T, and NUGC3), we found that in transformed cells, IQGAP3 staining exhibited a more punctate morphology and showed a higher nuclear presence compared to non-transformed cells (Fig. S6A-C).

### IQGAP3 knock-out decreases β-catenin levels in NUGC3 and AGS cells

The canonical Wnt pathway is best known for its regulation of cell proliferation (Jung & Park, 2020). We next ascertained how IQGAP3 status affects Wnt-driven proliferative potential, we performed CRISPR-mediated IQGAP3 knock-out in two gastric cancer cell lines: the *TP53* mutant NUGC3 and the constitutively active Tcf transcriptional activity (CTTA) AGS cell line (Caca, Kolligs et al., 1999). As screened previously, both cell lines exhibit high IQGAP3 expression (Fig. 7A), making them suitable candidates for CRISPR-mediated knock-out.

IQGAP3-KO NUGC3 cells showed reduced total β-catenin levels (Fig. 8A) and decreased expression of Wnt target genes (*CCND1* and *MYC*) (Fig. 8B) compared to NUGC3 wild-type cells. Moreover, both cell proliferation (Fig. 8C) and clonogenic growth (Fig. 8D and E) were also impaired.

**Fig. 8.**
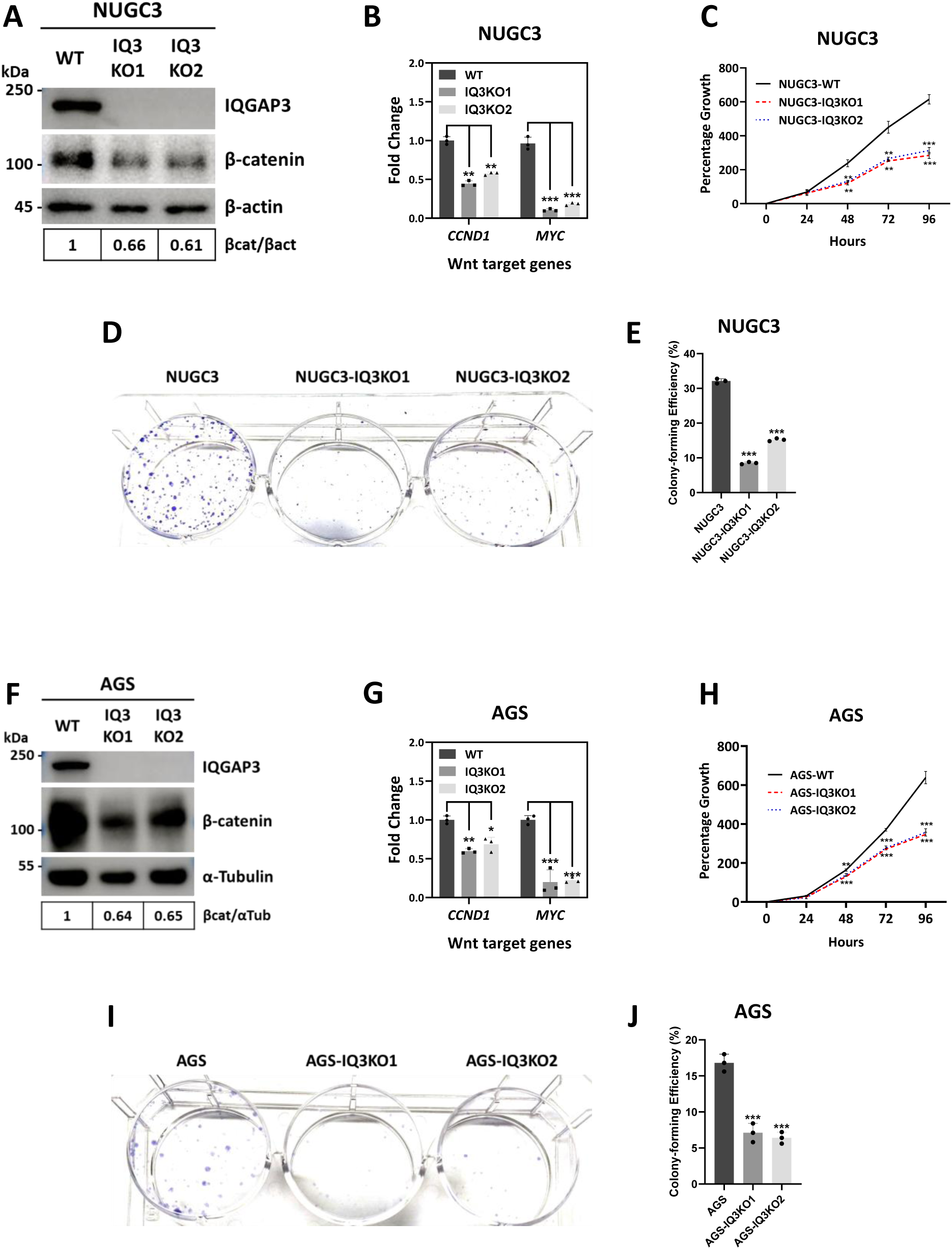
Loss of IQGAP3 reduces β-catenin levels in NUGC3 and AGS cells. (A) Immunoblot of NUGC3 wild-type and IQGAP3 CRISPR knock-out clones. Samples were subjected to SDS-PAGE (7.5%) followed by immunoblotting using the indicated antibodies. The relative immunoblot bands of β-catenin and β-actin (βcat/βact) were quantified by densitometry. (B) NUGC3 cells were lysed, and RNA collected were converted to cDNA and subjected to RT-PCR to quantify the amount of Wnt target genes *CCND1* and *MYC.* Representative data were collected and are expressed as the mean ± SD from three independent experiments. Student’s t test was performed, with ∗p < 0.05, ∗∗p < 0.01, and ∗∗∗p < 0.001. (C) XTT Cell Proliferation Assay of NUGC3 wild-type and IQGAP3 CRISPR knock-out clones. Representative data were collected and are expressed as the mean % growth ± SD from three independent experiments. Two-way ANOVA was performed, with ∗p < 0.05, ∗∗p < 0.01, and ∗∗∗p < 0.001. (D) Clonogenic Assay of NUGC3 wild-type and IQGAP3 CRISPR knock-out clones. (E) Colony-Forming Efficiency (%) of NUGC3 wild-type and IQGAP3 CRISPR knock-out clones. (F) Immunoblot of AGS wild-type and IQGAP3 CRISPR knock-out clones. Image is best representative of three independent experiments. Samples were subjected to SDS-PAGE (7.5%) followed by immunoblotting using the indicated antibodies. The relative immunoblot bands of β-catenin and α-Tubulin (βcat/αTub) were quantified by densitometry. (G) AGS cells were lysed, and RNA collected were converted to cDNA and subjected to RT-PCR to quantify the amount of Wnt target genes *CCND1* and *MYC.* Representative data were collected and are expressed as the mean ± SD from three independent experiments. Student’s t test was performed, with ∗p < 0.05, ∗∗p < 0.01, and ∗∗∗p < 0.001. (H) XTT Cell Proliferation Assay of AGS wild-type and IQGAP3 CRISPR knock-out clones. Representative data were collected and are expressed as the mean % growth ± SD from three independent experiments. Two-way ANOVA was performed, with ∗p < 0.05, ∗∗p < 0.01, and ∗∗∗p < 0.001. (I) Clonogenic Assay of AGS wild-type and IQGAP3 CRISPR knock-out clones. (J) Colony-Forming Efficiency (%) of AGS wild-type and IQGAP3 CRISPR knock-out clones.

Relevant to Wnt signaling, AGS harbor mutations for the following proteins; *CTNNB1* (G34E) (Caca et al., 1999) and *CDH1 (*G579fs) (Caldeira, Figueiredo et al., 2015). These mutations result in the accumulation of β-catenin and loss of E-cadherin expression, respectively. Loss of IQGAP3 impaired cell proliferation and clonogenic growth in the AGS gastric cancer cell line (Fig. 8H-J). Importantly, AGS IQGAP3 knock-out (IQ3KO) displayed significant loss of β-catenin relative to the AGS parental counterpart (Fig. 8F). Concomitantly, the expression of Wnt target genes involved in promoting proliferation and cell-cycle re-entry, *CCND1* and *Myc* (Niehrs & Acebron, 2012, Nusse, 2008, van de Wetering, Sancho et al., 2002) were also reduced in AGS IQ3KO cells (Fig. 8G).

In line with our findings, the relationship between IQGAP3 expression and β-catenin levels has also been previously observed, whereby the loss of IQGAP3 expression via siRNA-mediated knockdown in pancreatic cancer cell lines BxPC-3 and SW1990 have been shown to reduce β-catenin expression (Xu, Xu et al., 2016). Hence, our observation that the loss of IQGAP3 reduces β-catenin expression in gastric cancer cells may also apply to other cancer types.

### IQGAP3 is a target of Wnt signaling, sustains Wnt signaling via positive feedback loop

Wnt activity in epithelial stem cells are tightly regulated. Wnt-signaling levels below a specific threshold can be lethal due to the rapid turnover rate of the epithelium, as observed in mouse models of Tcf4 conditional knock-out (van Es, Haegebarth et al., 2012) or DKK overexpression (Kuhnert, Davis et al., 2004). Conversely, excessively high Wnt-signaling resulted in the overproduction of crypts, the formation of adenomas, and the development of cancer (Polakis, 2012). To maintain stringent control, the pathway incorporates various molecular checkpoints and feedback loops, where pathway components (e.g. AXIN2 (Leung, Kolligs et al., 2002), DKK1 (Niida, Hiroko et al., 2004), SFRP1 (Caldwell, Jones et al., 2006), WIF1 (He, Reguart et al., 2005), CD44 (Schmitt, Metzger et al., 2015), and NKD1 (Van Raay, Coffey et al., 2007)) play distinct regulatory roles.

To assess whether IQGAP3 is involved in the Wnt feedback loop, we treated HEK293T cells with Wnt3a conditioned media and analyzed IQGAP3 transcript and protein expression. As expected, RT-qPCR revealed increased expression of known Wnt targets, including AXIN2, CCND1, MYC, LEF1, TCF7, and TCF7L2 (also known as TCF4), following Wnt3a conditioned media treatment.

Interestingly, we also observed a Wnt3a-associated increase in IQGAP3 transcript levels (Fig. 9A). Supporting this, a time-course analysis of Wnt3a conditioned media treatment in HEK293T cells showed elevated IQGAP3 expression over time (Fig. 9B). Additionally, probing for IQGAP3 expression in L cells and L-Wnt3a cells revealed higher IQGAP3 levels in L-Wnt3a cells compared to L cells (Fig. S6D). Together, these findings strongly suggest that IQGAP3 is a Wnt target gene.

**Fig. 9.**
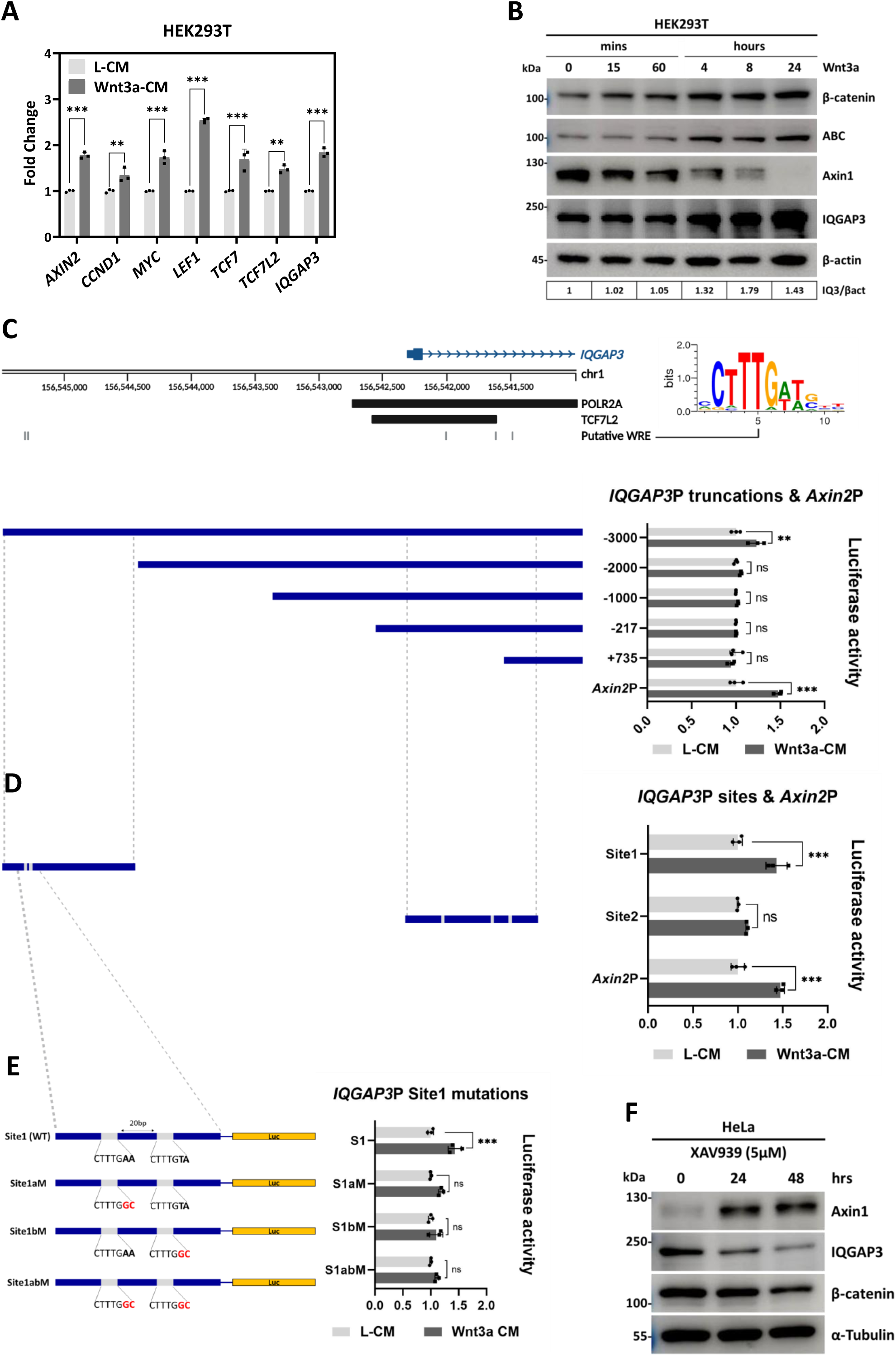
IQGAP3 is a target of Wnt signaling, sustains Wnt signaling via a positive feedback loop. (A) HEK293T cells were treated with wnt3a conditioned media supplemented with 100ng/ml of rhWnt3a. Cells were harvested, and the RNA collected were converted to cDNA and subjected to RT-PCR to quantify the amount of *AXIN2, CCND1, MYC, LEF1, TCF7, TCF7L2* and *IQGAP3* transcripts levels. (B) Time-course of HEK293T cells treated with Wnt3a conditioned media supplemented with 100ng/ml of rhWnt3a. Image is best representative of three independent experiments. Samples were subjected to SDS-PAGE (7.5%) followed by immunoblotting using the indicated antibodies. The relative immunoblot bands of IQGAP3 and β-actin (IQ3/βact) were quantified by densitometry. (C) Luciferase assay of 5’ truncation of IQGAP3 promoter region upon treatment with L-cells conditioned or wnt3a conditioned media. (D) Luciferase assay of Site1 and Site2 within the IQGAP3 promoter region upon treatment with L-cells conditioned or wnt3a conditioned media. (E) Luciferase assay of Site1, Site1a mutant, Site1b mutant and Site1ab mutant upon treatment with L-cells conditioned or wnt3a conditioned media. (F) Time-course of HeLa cells treated with 5µM of XAV939. Samples were subjected to SDS-PAGE (7.5%) followed by immunoblotting using the indicated antibodies. All the above data are representative of three independent experiments.

We next investigated how Wnt signaling regulates *IQGAP3* transcription. The Genome browser revealed multiple canonical TCF4 binding motifs (A/T-A/T-C-A-A-A-G or C-T-T-T-G-A/T-A/T) (van de Wetering & Clevers, 1992) within the *IQGAP3* promoter (*IQGAP3*P). The promoter region of *IQGAP3* spanning 4320bp (−3000 to +1320bp of TSS), a series of *IQGAP3*P 5’-truncations, and *Axin2* promoter *(Axin2*P) spanning 1730bp (−581 to +1149bp of TSS) were cloned into the pGL3-basic firefly luciferase reporter vector. We observed that both the full length *IQGAP3*P (−3000 to +1320 of TSS) and *Axin2*P responded to Wnt3a treatment, while the truncated *IQGAP3*P constructs did not (Fig. 9C). Thus, we hypothesized that a Wnt Responsive Element (WRE) might reside within -3000 to -2000bp of the TSS. Indeed, a truncation comprising the smaller fragment of -3000 to -2000bp (Site 1) displayed strong luciferase activity upon Wnt3a-CM treatment, similar to *Axin2*P, whereas negative control Site 2 (+335 to +1265 of the TSS) did not (Fig. 9D). Within Site1 we identified two potential WREs located within -2815 to -2808bp and -2810 to - 2803bp of the TSS, harboring the motifs CTTTGAA and CTTTGTA, respectively. Next, we mutated the motifs: CTTTG<u>AA</u> to CTTTG<u>GC</u> (Site1aM) and CTTTG<u>TA</u> to CTTTG<u>GC</u> (Site1bM) and observed that both Site1a and Site1b are necessary for Wnt3a-mediated upregulation of IQGAP3 transcription (Fig.9E). Lastly, we validated whether inhibition of Wnt signaling affects IQGAP3 expression by treating cells with XAV939, a Tankyrase (TNKS) inhibitor, that functions to stabilize Axin1 (Huang, Mishina et al., 2009). Time-course treatment with XAV939 increases Axin1 expression while decreasing both β-catenin and IQGAP3 expression (Fig 9F). Taken together, our findings confirm that IQGAP3 is a Wnt target.

## DISCUSSION

Scaffold protein IQGAP3 is upregulated in most, if not all, cancer cells and has been shown to be required for proliferation (Nojima et al., 2008). Identifying the IQGAP3 interactome in cancer cells is thus a critical step to develop anti-IQGAP3 therapeutics in cancer. Traditional immunoprecipitation/mass spectrometry techniques may miss transient, but crucial interactions in the IQGAP3 scaffold function. Here, we utilize Proximity ligation labelling to systemically characterize the molecular interactions and mechanisms underlying the IQGAP3 scaffold function in cancer.

We identified large networks of IQGAP3 interacting proteins in the cell cycle as well as oncogenic pathways. Based on our interactome data, IQGAP3 is associated with different protein complexes located at various locations of the cell and phases of the cell cycle. We observed that IQGAP3 proximity partners were enriched for proteins found in non-membrane bounded organelle or phase separated microenvironment. By forming interactions with other phase-separated scaffold proteins, IQGAP3 could penetrate the specialized microenvironments of various biochemical reactions and thus regulate these reactions.

In this study, we have uncovered a previously unknown function of IQGAP3. IQGAP3 expression disrupts Axin1-CK1α interaction, either by blocking CK1α binding to Axin1, sequestering CK1α away from Axin1, or both, thereby inhibiting β-catenin phosphorylation and promoting its stabilization (Fig 10). Notably, we identified that the coiled-coil (CC) domain of IQGAP3 is essential for its phase separation, while the IQ domain is necessary for its nuclear localization. Both functions are critical for IQGAP3 to enhance β-catenin activity. This dependence on the coiled-coil domain to phase separate resembles proteins such as, Amot (Wang, Choi et al., 2022, Wells, Fawcett et al., 2006), KIBRA (Wang et al., 2022), SLMAP (Wang et al., 2022), TAZ (Lu, Wu et al., 2020), YAP (Mao, Liu et al., 2023), liprin-α (Liang, Jin et al., 2021), LINE-1 ORF (Newton, Naik et al., 2021), and RNA-Binding Proteins (RBPs) (Ford & Fioriti, 2020). In addition, the IQ domain potentially mediates IQGAP3 nuclear localization through hitchhiking on proteins harboring an NLS. Elucidating this mechanism could provide insight into the role of IQGAP3 in the nucleus.

**Fig. 10:**
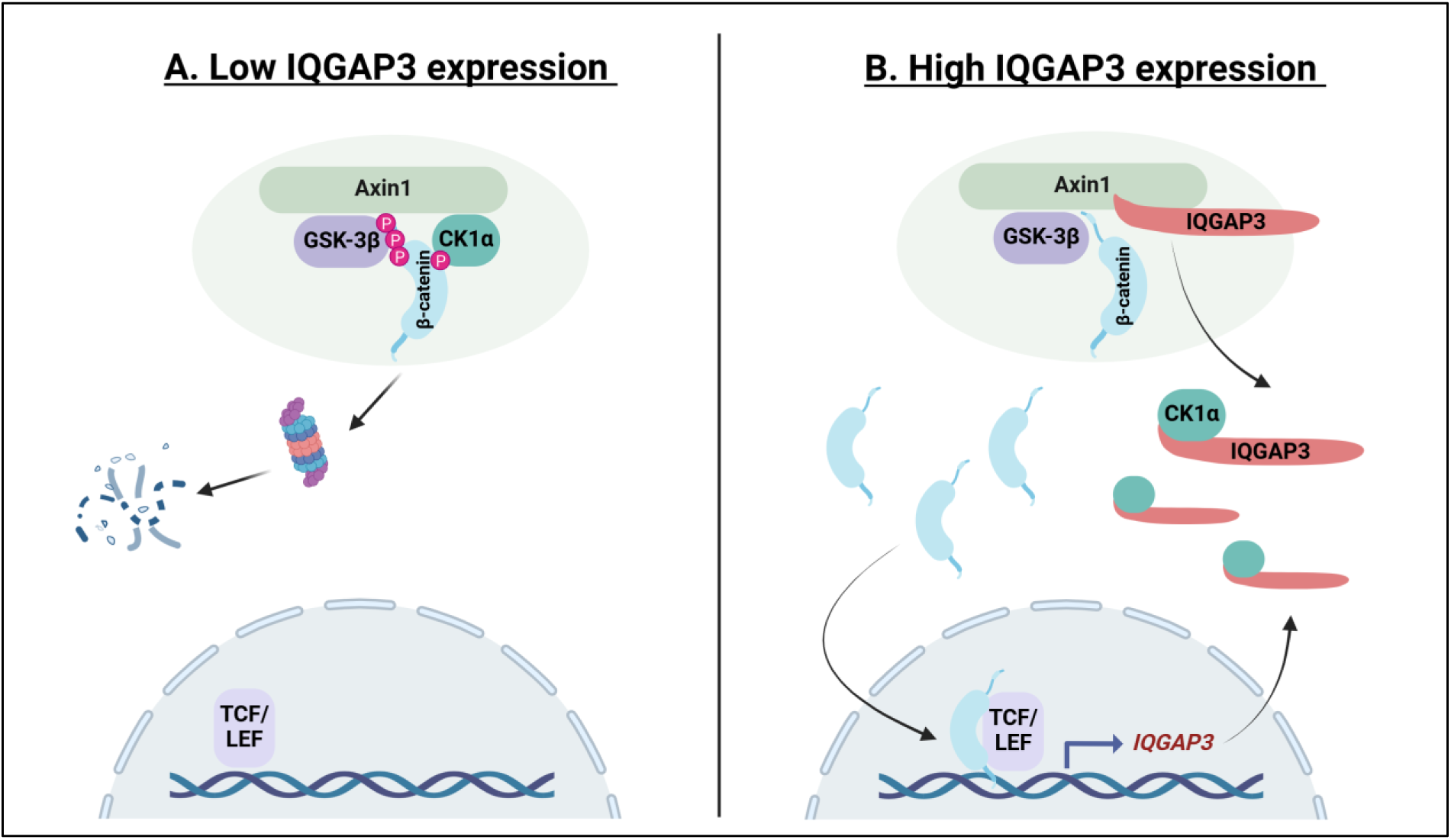
Proposed model for IQGAP3 role in Wnt signalling. We proposed that In IQGAP3-low cells (A), the destruction complex phosphorylates β-catenin, targeting it for degradation. Meanwhile, in IQGAP3-high cells (B), IQGAP3 competes with CK1α for binding to Axin1 or sequesters it away from Axin1 or both, thus, inhibiting CK1α-mediated β-catenin phosphorylation. This inhibition leads to the accumulation of β-catenin, which promotes the expression of Wnt target genes, including IQGAP3, Created with BioRender.com.

In addition, we observed that Wnt3a treatment promotes IQGAP3 expression both at the transcript and protein levels. Wnt signaling is tightly regulated with multiple feedback loops, where most Wnt regulators are themselves Wnt target genes (Zhong & Virshup, 2020). Similarly, IQGAP3 is both a regulator and target of Wnt signaling. This Wnt-associated function of IQGAP3 supports previous findings that implicate IQGAP3 in promotion of cancer proliferation and stem cell regeneration. Therefore, targeting IQGAP3 during early stages of cancer could potentially help mitigate both the growth and spread of cancers.

## MATERIALS AND METHODS

### Cell culture, transfection and conditioned media

HEK293T (CRL-3216), L-Cells (CRL-2648), and L-Wnt3a cells (CRL-2647) were from ATCC. HEK293 Tet-On® 3G Cell Line #631182 was from Takara bio. Immortalized human normal gastric epithelial cells (HFE145) were obtained from Dr. Hassan Ashktorab and Dr. Duane Smoot. Both HEK293 and HEK293T cells were cultured in Dulbecco’s modified Eagle’s medium (DMEM) while the gastric cancer cell lines, AGS, MKN1, MKN28, HFE145 and NUGC3 cells were cultured in RPMI-1640, both growth media were supplemented with 10% fetal bovine serum (FBS) and 1% Penicillin-Streptomycin solution. The Dox-inducible 3×HA-TurboID-empty and 3×HA-TurboID-IQGAP3 expressing cells were generated by transfection of pRetroX-3×HA-TurboID-empty or pRetroX-3×HA-TurboID-IQGAP3 in the HEK293-Tet-On and MKN28-Tet-On cells, thereafter, stable population were selected with 1.0 µg/ml Puromycin.

All Dox-inducible TurboID stable expressing cells were cultured and maintained in DMEM supplemented with 10% fetal bovine serum (FBS), 0.5 µg/ml Puromycin (Gibco), 1 µg/ml Doxycycline (Sigma), 200 µg/ml Geneticin (Gibco). Transfections were carried out using Lipofectamine 2000 (Thermofisher) according to the manufacturer’s instructions. CRISPR knock-out cells were cultured and maintained in RPMI supplemented with 10% fetal bovine serum (FBS), and 0.5 µg/ml Puromycin. Conditioned media from L-cells and L-Wnt3a cells were harvested 72-96 hours after cell seeding and filtered through 0.45 µm syringe filter and stored in -80°C. All Wnt3a conditioned media were supplemented with 100 ng/ml rhWnt3a #5036-WN (RnDSystems).

All cell lines were verified for authenticity prior to use and screened for mycoplasma contamination every three months using the LookOut® Mycoplasma qPCR Detection Kit (Sigma Aldrich).

### Antibodies and Plasmids

Antibodies specific for the following proteins were used at the indicated dilutions for Immunoblot: α-Tubulin 11224-1-AP (1:10,000, Proteintech®); IQGAP3 #25930-1-AP, mCherry #26765-1-AP (all 1:1000, Proteintech®); Flag M2 #F1804 (1:1000, Sigma); Axin1 #2087, β-catenin #8480, Non-phospho (Active) β-Catenin (Ser45) #19807, Non-phospho (Active) β-Catenin (Ser33/37/Thr41) #8814, Myc-Tag #2276, HA-Tag #3724 (rabbit) & #2367 (mouse), GSK-3β #12456 (rabbit) & #9832 (mouse), PP2A C #2038 (all 1:1000, Cell Signaling). Streptavidin-HRP #ab7403 (1:40,000, Abcam); IQGAP1 #ab86064 (1:1000, Abcam), CK1α Antibody (H-7): sc-74582 (1:250, Santa Cruz).

Antibodies specific for the following proteins were used at the indicated dilutions for immunoprecipitation: Axin1 #2087 (1:100, Cell Signaling), Myc-Tag (9B11) #2276 (1:100, Cell Signaling). Anti-Flag® M2 #F1804 (1:100 Sigma), IQGAP3 (H00128239PW2) #89-301-822 (1:100, Abnova).

The following plasmid was obtained from Vectorbuilder:

pCMV-EGFP-IQGAP3

The following plasmids were obtained from Addgene:

3xHA-TurboID pRetrox (from Kyle Roux, #155203), pGL3-OT and pGL3-OF (from Bert Vogelstein, #16558 & #16559), Flag-Axin1 (from Mariann Bienz, #109370), β-catenin-Flag (from Eric Fearon, #16828), LentiCRISPRv2 (from Feng Zheng, #52961), pMD2.G and psPAX2 (from Didier Trono #12259 and #12260), mCherry2-C1 (from Michael Davidson, #54563), pcDNA3-EGFP-empty (Doug Golenbock, #13031), pcDNA3-GSK3β-HA (from Jim Woodgett, # 14753), pLC-Flag-CSNK1A1 WT-Puro (from Eva Gottwein #123319)

### Molecular cloning

Cloning was performed using either In-Fusion HD Cloning Kits (Takara) via Ligation Independent Cloning (LIC) or Restriction cloning using enzymes purchased from New England Biolabs (NEB).

Full-length IQGAP3 coding sequence was cloned from pCMV-IQGAP3-GFP and inserted into 3xHA-TurboID pRetrox, pcDNA3-Flag-empty, pcDNA3-Myc-empty, and mCherry2-C1. Full-length Axin1 coding sequence was cloned from Flag-Axin1 and inserted into pcDNA3-EGFP-empty and pcDNA3-HA-empty. Full-length β-catenin coding sequence was cloned from β-catenin-Flag and inserted into pcDNA3-HA-empty. Full-length CK1α coding sequence was cloned from pLC-Flag-CSNK1A1 WT-Puro and inserted into pcDNA3-HA-empty.

cDNA expressing EGFP-CC was amplified from the pCMV-EGFP-IQGAP3 vector using the following primers: EcoRI-CC-For, TAAGCAGAATTCCTCTTCCTCTTCCGGCTGGG and XhoI-CC-Rev, TGCTTACTCGAGGAAAGCTGTGTCTGCTGG generating a 1593 bp primer product that was later cloned into an EGFP-vector backbone. Similarly, EGFP-IQ was amplified from pCMV-EGFP-IQGAP3 vector using the following primers: EcoRI-IQ-For, TAAGCAGAATTCGTTATCCAGCTCCAGGCCCG and XhoI-IQ-Rev, TGCTTACTCGAGCTAAGGGTGGGGTGCATGCAC generating a 375 bp primer product that was later cloned into an EGFP-vector backbone.

### Promoter cloning

Promoter sequences of *AXIN2* and *IQGAP3* were obtained from Ensembl. *IQGAP3* promoter (*IQGAP3P*): A region spanning 4320bp located on chr1: 156,571,285-156,575,604 (GRCh38/hg38) or chr1:156,541,077-156,545,396 (GRCh37/hg19) and *AXIN2* promoter (*AXIN2P*): A region spanning 1730bp located on chr17: 65,560,500-65,562,229 (GRCh38/hg38) or chr17:63,556,618-63,558,347 (GRCh37/hg19). Both promoter sequences were PCR amplified using genomic DNA isolated from HEK293T cells with KAPA HiFi PCR Kit. Primer-BLAST was used to verify the specificity of primer design. In-Fusion HD Cloning Kits (Takara) were used in Ligation Independent Cloning (LIC) of PCR product into pGL3 basic vector. Primer sequences excluding homology sequences (added for LIC) are as follows, for *IQGAP3P* forward primer: 5’-ACCTATAGATGTAACTTGTCCAAG-’3 and reverse primer: 5’-GTAAGAAAATAAACTCAGAGAGCTT-3’, for *AXIN2P* forward primer: 5’-CACTCAGGCCTATACTGGCG-3’ and reverse primer: 5’-CACGCCGACTCACATCCATA-3’. All promoter plasmid constructs were co-transfected with pSV-β-Galactosidase (internal control) into HEK293T cells with and without Wnt3a-CM treatment to validate the putative Wnt responsive element (WRE) within the promoter region.

### gRNA design and plasmids

The sgRNA sequence including PAM region targeting exon 5 of *IQGAP3* was determined using IDT (sg.idtdna.com/site/order/designtool/index/CRISPR_SEQUENCE) as follows; 5’-TGGATGCAGTAGACTACCCG GGG-3’. While sgRNA targeting exon 17 of *IQGAP3* was determined using CRISPR Design Tool (www.genscript.com/CRISPR-gRNA-constructs.html) as follows; 5’-GTCGGGAACTACCCCTCGAA GGG-3’. sgRNA oligos were cloned into LentiCRISPRv2 plasmid (Addgene #52961) at BsmBI restriction sites (Sanjana, Shalem et al., 2014).

### Lentiviral vector construction and transduction

Lentiviral vectors were produced by co-transfecting HEK293T cells with the transfer plasmid sgRNA-Cas9-expressing lentiCRISPRv2, the packaging plasmid pMD2.G and psPAX2 using Lipofectamine 2000 transfection reagent. Briefly, for each sgRNA, a plate of 10-cm plate of 60% confluent 293T cells was transfected with 10 μg of the transfer plasmid, 5μg of pMD2.G, 7.5μg of psPAX2, and 40 ul of Lipofectamine 2000 (Life Technologies). After 24 hours, the media was replaced with fresh media. After 48 hours, viral supernatants were collected and centrifuged at 3,000 rpm at 4 °C for 5 min to pellet cell debris. The supernatant was filtered through a 0.45 μm low protein binding membrane (Millipore) and used immediately after supplementing with 7μg/ml of polybrene or stored in −80°C. Successful knock-out colonies were selected with Puromycin treatment, 1 μg /ml.

### Immunoblot

Cells were lysed with RIPA lysis buffer (ThermoSci) supplemented with Protease inhibitor cocktail (Roche) and Halt™ Phosphatase Inhibitor Cocktail (Thermo Scientific™). For all unless specified, 20 µg of total protein from each sample were subjected to 7.5% or 10% SDS polyacrylamide gel electrophoresis (PAGE) and electrophoretic transfer onto 0.2 µm polyvinylidene difluoride (PVDF) membrane (Bio-Rad) at 30 V, 100 mA, 4 °C overnight. The transfer buffer (Towbin) comprises 25 mM Tris, 192 mM glycine, 20% (v/v) methanol, pH 8.3. Membranes were incubated with primary antibodies in either 0.5% BSA or skimmed milk in TBST (Tris Buffered Saline + 0.1% (v/v) Tween 20) overnight, following which were washed 3 times with TBST and incubated with secondary antibody conjugated with HRP (1:5000) in 0.5% BSA or skimmed milk in TBST for 1 hour at room temperature. Membranes were then washed 3 times with TBST. Antibody detection was performed using Radiance plus chemiluminescent substrate (azure biosystems). Antibody reactions detection was imaged using ImageQuant LAS 500 (GE Healthcare). When re-probing with a different primary antibody, membranes were stripped using Mild stripping buffer pH 2.2 (1.5% (w/v) Glycine, 1% (w/v) SDS, and 1% Tween 20) in room temperature for 10 minutes and thereafter washed with TBS for 10 minutes twice and with TBST for 10 minutes twice. All immunoblotting were performed thrice, only best represented images were used. Uncropped images of all immunoblots can be found in the supplementary image file.

### Immunofluorescence staining

Cells were fixed with 4% paraformaldehyde for 30 minutes and washed with PBS three times for 5 minutes each. Cells were then permeabilized with PBS containing 0.1% Triton X-100 for 10 minutes. Permeabilized cells were then incubated in blocking buffer (0.5% BSA/0.1% Triton X-100) for 1 hour, and following that with the appropriate primary antibody diluted in antibody dilution buffer (0.5% BSA/0.3% Triton X-100) overnight in 4°C. After three washes with PBS for 5 minutes each, cells were incubated with Alexa Fluor 488- or 594-conjugated secondary antibodies (1:100 dilution) for 1 hour. Cells were then washed with PBS three times for 5 minutes each and mounted with DAPI. Fluorescence was observed under Zeiss LSM 880 Airy Scan.

### Live Cell Imaging

HeLa Tet-On cells stably expressing Doxycycline inducible EGFP-empty and EGFP-IQGAP3 were seeded onto a 35 mm cell imaging dish with a glass bottom (Matsunami). Cells were induced with Doxycycline 24-48 hours prior to live imaging.

### Fluorescence recovery after photobleaching (FRAP)

HeLa cells grown on Glass Bottom Culture Dishes (Matsunami Glass) were transfected with 2 μg per well of GFP–CC and analyzed through quantitative FRAP studies 24h after transfection. FRAP experiments were performed using a Zeiss LSM880 confocal microscope. The fluorescence signal was bleached using 100% of maximum laser power of the 488-nm laser with 150 interactions for ∼2.5 s. Relative recovery was normalized to the initial (before bleaching) fluorescence intensity for each bleached condensate and then used to calculate mean and SD of the recovery time. To compare the post-bleach recovery 5 pre-bleach images were captured.

For the treatment of 1,6-hexanediol (1,6-HD), culture media was replaced with 5% 1,6 HD prepared in DMEM culture media, which was replaced with fresh culture media to monitor the recovery after 1,6 HD treatment. Apart from the bleached region, fluorescence intensity mean was also recorded for non-bleached and background. Raw FRAP data was further analyzed using easyFRAP online analysis tool (Koulouras et al., 2018). All acquired images were processed and analyzed in ImageJ software.

### Co-immunoprecipitation

Cells were lysed with immunoprecipitation lysis buffer (IPLB) containing 25 mM Tris pH 7.5, 150 mM NaCl, 0.5% Triton-X, protease inhibitor cocktail (Roche) and Halt™ Phosphatase Inhibitor Cocktail (Thermo Scientific™). For Flag-tagged and endogenous protein immunoprecipitation, cell lysates were incubated with Protein G magnetic beads (Millipore) for 2 hours (pre-clearing). The beads were removed, and primary antibody was then added to protein lysates at a concentration of 1:100 and incubated overnight at 4°C. 15 µl of Protein G magnetic beads (Millipore) is then added to the Protein-Antibody complex and incubated at 4°C for 2 hours. The Protein G magnetic beads (Millipore) were then washed with IPLB three times and protein complex were eluted by adding SDS sample buffer and boiling for 5 minutes.

### Biotin pulldowns

Using the TurboID method (Branon et al., 2018), proteins in close proximity to IQGAP3 were biotinylated and isolated by streptavidin-bead pulldowns. Briefly, 24 hr after Dox induction and biotin supplementation (50 μM), biotinylated proteins were enriched from the protein extracts with 50 µL of streptavidin-coated magnetic beads (Dynabeads™ MyOne™ Streptavidin C1, Invitrogen, #65001). Beads were washed twice with RIPA lysis buffer, the lysates containing 2 mg protein were then incubated with the equilibrated beads on a rotator overnight at 4 °C. The beads were washed twice each with wash buffer 1 (2% SDS), wash buffer 2 (50 mM Hepes pH 7.5, 500 mM NaCl, 1 mM EDTA, 1% Triton-X-100, 0.5% Na-deoxycholate), wash buffer 3 (10 mM Tris pH 8, 250 mM LiCl, 1 mM EDTA, 0.5% NP-40, 0.5% Na-deoxycholate) and wash buffer 4 (50 mM Tris pH 7.5, 50 mM NaCl, 0.1% NP-40). Biotinylated proteins were eluted by boiling the beads in 35 μL of 1 × Laemmli sample buffer. Eluted proteins were sent for Mass spectrometry analysis.

### Mass spectrometry analysis

TurboID samples were separated on a 4-12% NuPAGE Bis-Tris precast gel (Thermo Fisher Scientific) for 10 min at 170 V in 1x MOPS buffer. The gel was fixed using the Colloidal Blue Staining Kit (Thermo Fisher Scientific) and processed as a single fraction. For in-gel digestion, samples were destained in destaining buffer (25 mM ammonium bicarbonate; 50% ethanol), reduced in 10 mM DTT for 1h at 56°C followed by alkylation with 55mM iodoacetamide (Sigma) for 45 min in the dark. Tryptic digest was performed in 50 mM ammonium bicarbonate buffer with 2 μg trypsin (Promega) at 37°C overnight. Peptides were desalted on StageTips and analysed by nanoflow liquid chromatography on an EASY-nLC 1200 system coupled to a Q Exactive HF mass spectrometer (Thermo Fisher Scientific). Peptides were separated on a C18-reversed phase column (25 cm long, 75 μm inner diameter) packed in-house with ReproSil-Pur C18-AQ 1.9 μm resin (Dr Maisch). The column was mounted on an Easy Flex Nano Source and temperature controlled by a column oven (Sonation) at 40°C. A 105-min gradient from 2 to 40% solution B (80% acetonitrile, 0.1% formic acid) at a flow of 225 nl/min was used. Spray voltage was set to 2.2 kV. The Q Exactive HF was operated with a TOP20 MS/MS spectra acquisition method per MS full scan. MS scans were conducted with 60,000 at a maximum injection time of 20 ms and MS/MS scans with 15,000 resolution at a maximum injection time of 75 ms. The raw files were processed with MaxQuant version 2.0.1.0 (HEK293) and 1.5.2.8. (MKN28) with preset standard settings for label-free quantitation using the MaxLFQ algorithm. Carbamidomethylation was set as fixed modification while methionine oxidation and protein N-acetylation were considered as variable modifications. Search results were filtered with a false discovery rate of 0.01. Known contaminants, proteins groups only identified by site, and reverse hits of the MaxQuant results were removed and only proteins with LFQ intensities in four replicates of at least one condition of each pair-wise comparison were kept. Missing LFQ intensities were randomly inputed from a normal distribution around the lowest 5% of LFQ intensities based on the means of three iterations.

### Cell Proliferation Assay

Cell proliferation was determined using the TACS® XTT Cell Proliferation Assay (R&D Systems, Inc.), following the manufacturer’s instructions. Gastric cancer cell lines NUGC3 and AGS along with their corresponding IQGAP3KOs were seeded at 1 × 10³ cells/mL density in 100 μL of appropriate growth medium in 96-well microplates. After incubation overnight, to allow cell attachment, which was considered as 0H and analyzed, followed by analysis after time periods of 24, 48, 72 and 96 hours. For analysis, 50 μL of XTT Working Solution (prepared by combining XTT Reagent and XTT Activator as per kit instructions) were added to each well. Plates were incubated for 6 hours at 37°C in a 5% CO₂ atmosphere. The absorbance was measured at 490 nm (test wavelength) with a reference wavelength of 630-690 nm using a microplate reader. The calculations are as follows: Cell Viability (%) = (A_t_ / A_0h_) × 100

Where: A_t_ = Absorbance of treated wells at respective time points (0, 24, 48, 72, 96 hrs) and A_0h_ = Absorbance of control wells at 0h.

### Clonogenic Assay

Gastric cancer cell lines NUGC3 and AGS along with their corresponding IQGAP3KOs were seeded in 6-well plates at a density of 500 cells per well in 2 mL of complete growth medium and incubated at 37°C in 5% CO₂. The cells were grown with necessary media change when required for 7–14 days to allow colony formation. At the end of the incubation period, the medium was removed, and cells were gently rinsed with PBS. Colonies were fixed and stained with a mixture of 6.0% glutaraldehyde and 0.5% crystal violet for 30 minutes. The staining solution was removed, and plates were rinsed in tap water and air-dried at room temperature. Colonies consisting of ≥50 cells were counted manually under a light microscope. The colony-forming efficiency (CFE) was calculated as: CFE (%) = Colonies formed / Cells seeded×100

### Luciferase assay

5 × 10^4^ HEK293T cells were grown per well in a 24-well plate for 24 hours, thereafter cells were transfected with the appropriate plasmids: for TOPFLash (TCF wild-type (OT) or TCF mutant (OF) plasmids); for promoter analysis (*IQ3P* or *Axin2P*), together with pSV-β-Galactosidase vector as control vector for monitoring the transfection efficiency. Cells were grown for an additional 48-hours before lysis. If Wnt3a conditioned media were used, cells were exposed to the conditioned media 24-hours before lysis. Lysis was performed using 5×Passive lysis buffer (Promega, #E1910), lysates were rocked in room temperature for 15 mins. Thereafter, lysates were centrifuged for 10 mins at 14,000 rpm to remove cell debris. Lysate supernatants used were aliquoted into white, flat bottom, 96-well plate (Costar) for treatment with either Beta-Glo® (Promega, #E4720) or Luciferase Assay Reagent II/LARII (Promega, #E1910) reagent.

### Real-Time Polymerase chain reaction (RT-PCR)

RNA extraction was performed using RNeasy Mini Kit (Qiagen). 1 µg of Purified RNA was reverse transcribed to complementary DNA (cDNA) using iScript™ Reverse Transcription Supermix (Bio-Rad). cDNA was diluted 10×. RT-PCR was performed on QuantStudio™ 3 Real-Time PCR System (Applied Biosystems). The 20 μl PCR reaction mixture consisted of 2 μl of cDNA, 100 nM forward and reverse primers, 2× iTaq Universal SYBR Green Supermix. Cycling conditions were initial denaturation at 95 °C for 5 minutes, followed by 40 cycles consisting of 95 °C for 10 seconds, 60 °C for 30 seconds and 72 °C for 10 seconds.

The primers used for real-time RT-PCR were as follows:

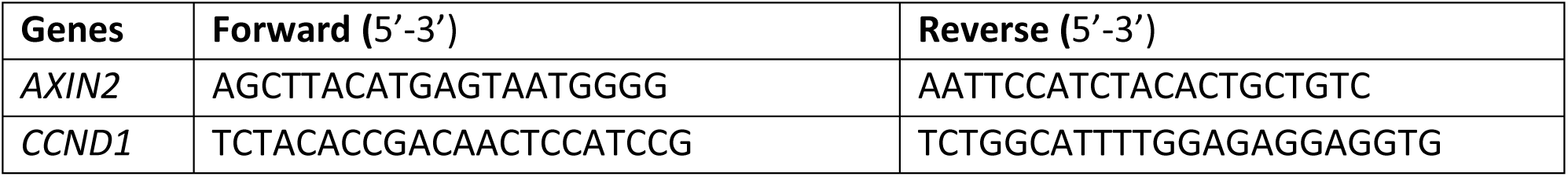
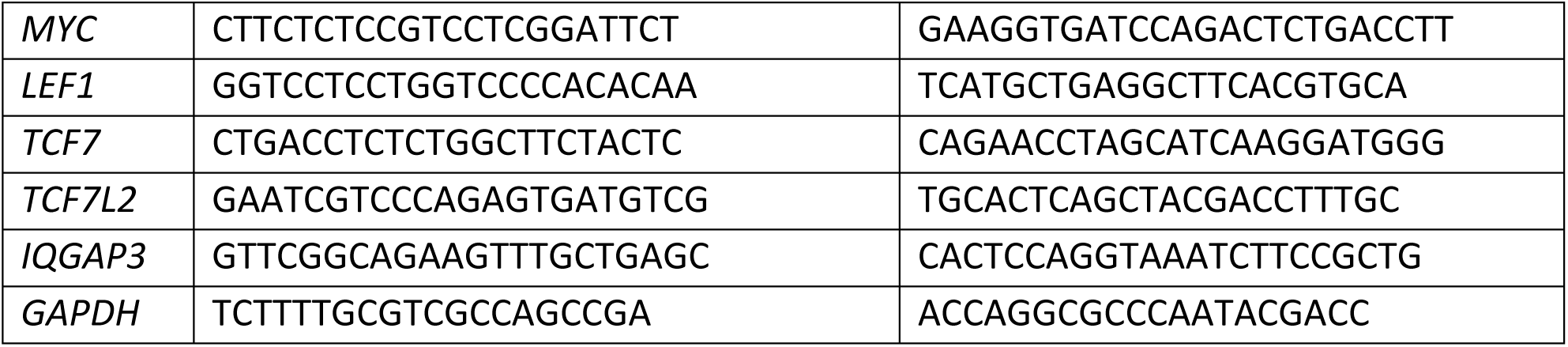

### Fluorescence Intensity Analysis

To determine mCherry2 intensity for localization of IQGAP1 and IQGAP3. The cross-sectional line was drawn in ImageJ. Lines were fixed by the ROI manager to ensure a similar length for image comparison. Lines were then analyzed using Plot Profile.

### Co-localization Analysis

To determine mCherry2/EGFP co-localization, channels were split in ImageJ and the JACoP plugin was used. The Pearson’s Coefficient generated from 50 cells were averaged.

### General Statistical Analysis

Experimental data are presented as the mean ± SD. Two-tailed Student’s t tests with unequal variance were performed using XLMiner Analysis ToolPak on Microsoft excel. Error bars represent standard deviation. Differences were considered statistically significant when p values were less than 0.05 (^∗^p < 0.05, ^∗∗^p < 0.01, ^∗∗∗^p < 0.001). The best represented images were shown.

## Supporting information

MassSpecData_HEK293_IQGAP3 -Wnt3a vs. Empty vector -Wnt3a

MassSpecData_HEK293_IQGAP3 +Wnt3a vs. Empty vector 377:59Wnt3a

MassSpecData_MNK28_IQGAP3_TurboID

RNAseq_4486_merged_gene_counts

Fig.3D_1.6HD_5mins

Fig.3D_2.5HD_5mins

Fig.3E_FRAP video_EGFP-IQGAP3

Fig.3E_FRAP_video_EGFP-IQGAP3 (FRAP & Fusion)

Fig.6H_FRAP video_EGFP-CC

## Author contributions

M.B.R. conceptualized the study. M.B.R. and Y.I. analyzed the data. A.H. performed XTT and Clonogenic assays. D.K. and S.P. performed the mass spectrometry analysis. M.B.R. and T.Y.X. performed the confocal imaging. M.B.R and B.D. performed the FRAP analysis. M.B.R. created the Tet-On and CRISPR-KO cells. M.B.R. and Y.I. wrote the manuscript.

## ACKNOWLEDGEMENTS

We thank Dr. Hassan Ashktorab and Dr. Duane Smoot for the Immortalized human normal gastric epithelial cells (HFE145). This research was supported by grants from the National Research Foundation Singapore and the Singapore Ministry of Education under its Research Centers of Excellence initiative, the Singapore Ministry of Health’s National Medical Research Council under its Clinician-Scientist Individual Research Grant (MOH-000933-00) and the National Medical Research Council’s Open Fund Large Collaborative Grant (OFLCG18May-0023), National University of Singapore School of Medicine (NUSMed) Internal Grant Funding (NUHSRO/2019/086/ StomachStemCell and NUHSRO/2022/043/NUSMed/25/LOA) to Y.I.

