## Supplementary Figures for "Wnt target IQGAP3 promotes Wnt signaling via disrupting Axin1-CK1α interaction"

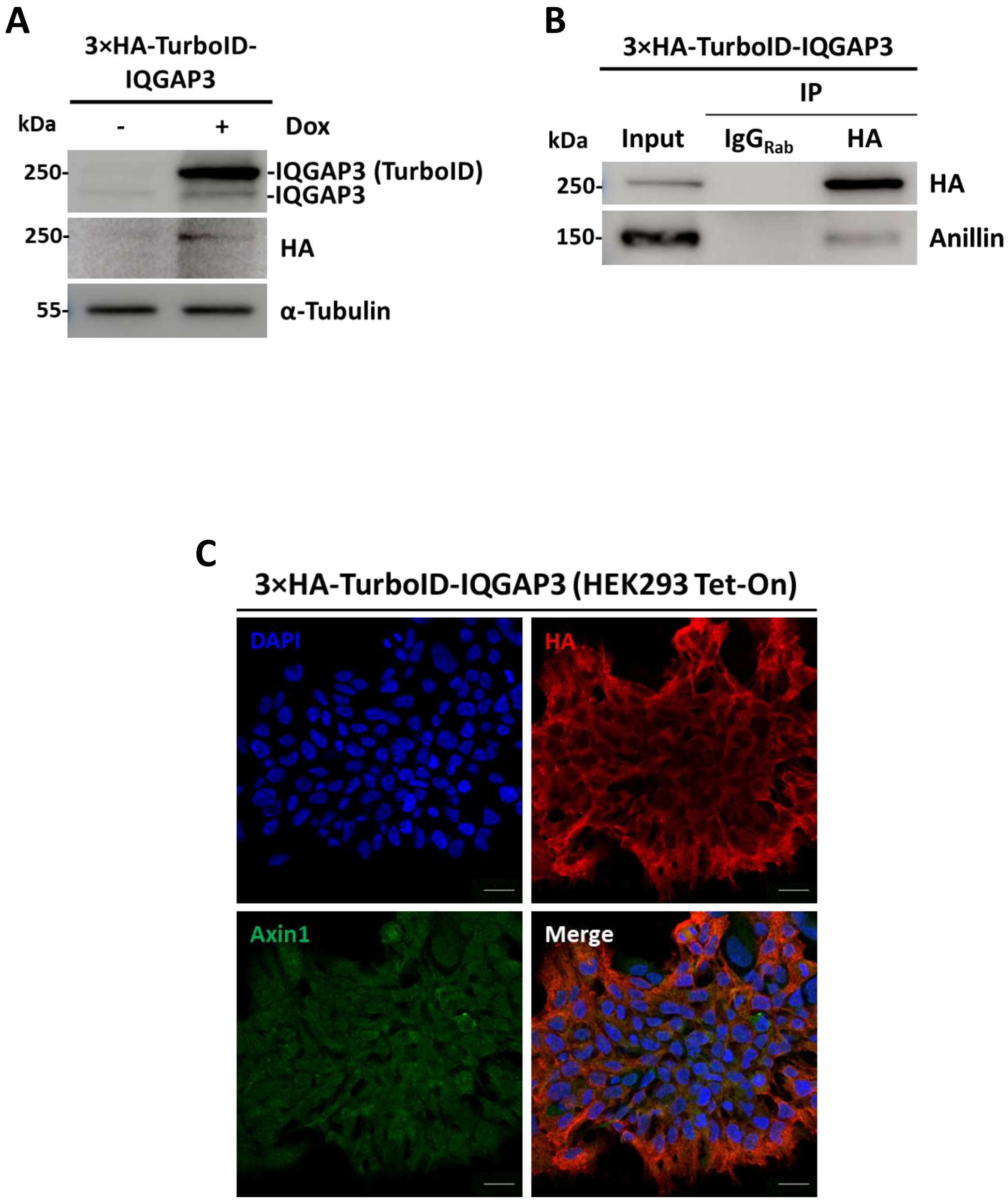
A
3×HA-TurboID-IQGAP3
kDa
-
+
Dox
250-
-IQGAP3 (TurboID)
-IQGAP3
250-
HA
55-
α-Tubulin
B
3×HA-TurboID-IQGAP3
IP
kDa
Input
IgG Rab
HA
250-
HA
150-
Anillin
C
3×HA-TurboID-IQGAP3 (HEK293 Tet-On)
DAPI
HA
Axin1
Merge

**Fig. S1: HEK293 3×HA-TurboID-IQGAP3 expression, immunoprecipitation and immunofluorescence staining.** (A) Doxycycline induction of TurboID-IQGAP3 in HEK293 cells probed with HA-tag (1:1000), IQGAP3 (1:1000), and α-tubulin (1:10,000) antibodies. (B) HA-tag Immunoprecipitation of lysates expressing 3×HA-TurboID-IQGAP3, probed for Anillin (1:1000) co-immunoprecipitation. (C) Immunofluorescence staining of HA-tag and Axin1 in cells expressing 3×HA-TurboID-IQGAP3, scale bars, 50µm.

A

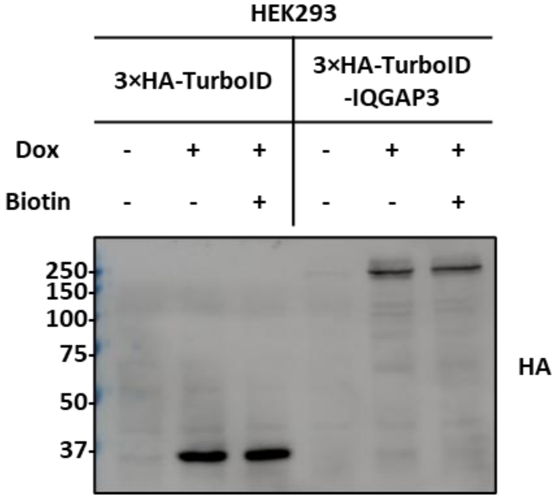


B

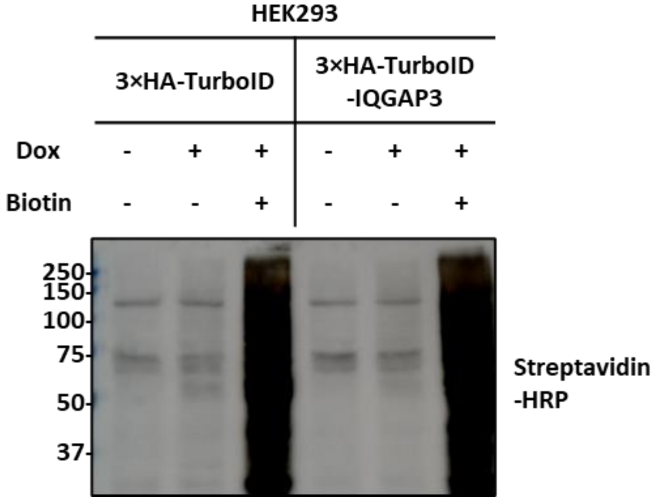


C

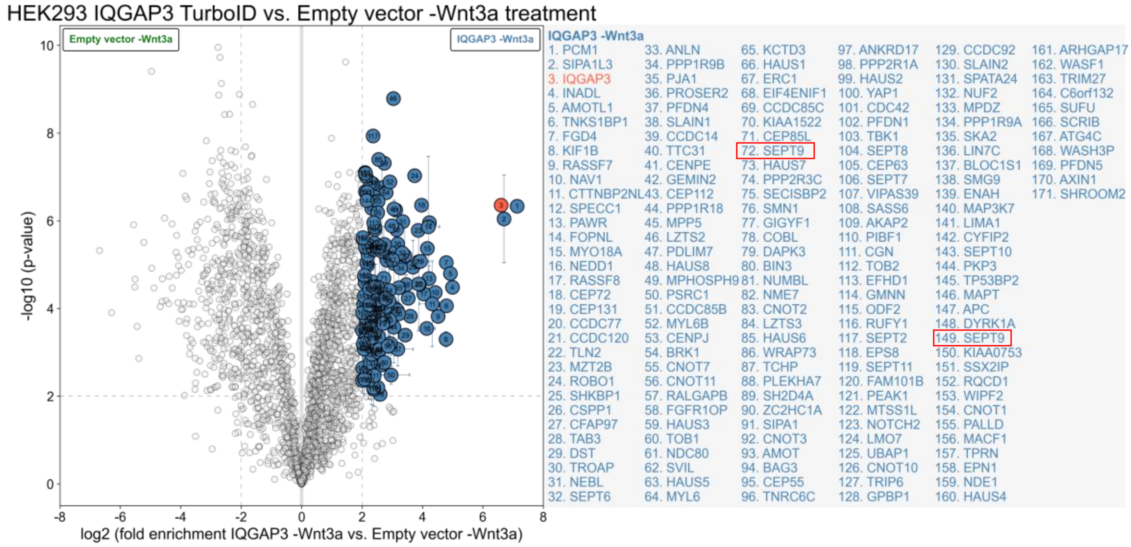


***SEPT9* has two different isoforms, hence appears twice in the Mass spectrometry analysis (#72 and #149).**

D

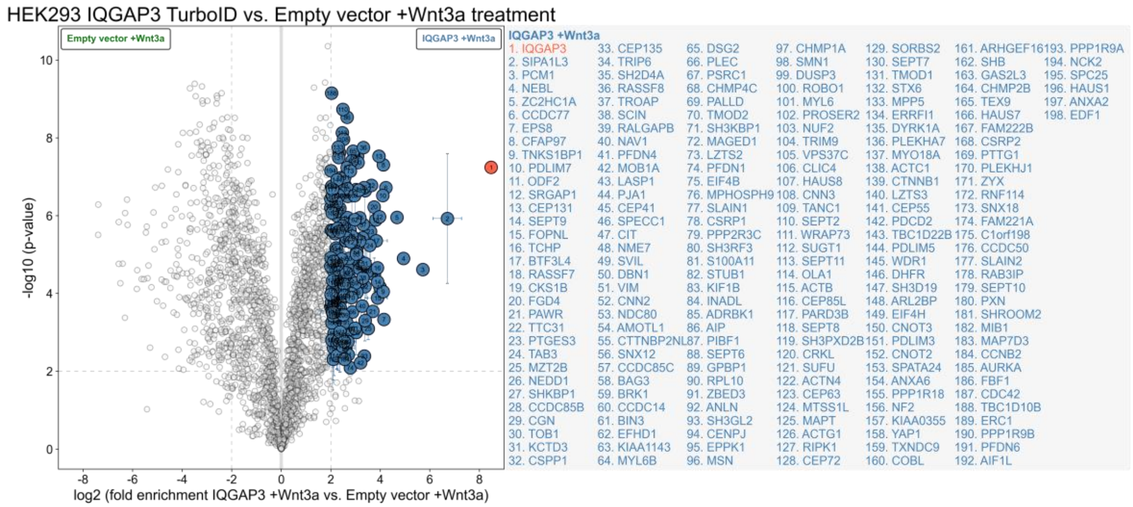

**Fig. S2: HEK293 TurboID-IQGAP3 induction, biotinylation and volcano plots.** (A) Doxycycline induction of TurboID-IQGAP3 in HEK293 cells probed with anti-HA antibody (1:1000). (B) Biotin labelling of TurboID-IQGAP3 in HEK293 cells probed with streptavidin-HRP antibody (1:40,000). (C) Volcano plot of TurboID-empty vs. TurboID-IQGAP3 (-Wnt3a) (Fold change ≥ 4, p-value ≤ 0.01). (D) Volcano plot of TurboID-empty vs. TurboID-IQGAP3 (+Wnt3a) (Fold change ≥ 4, p-value ≤ 0.01).

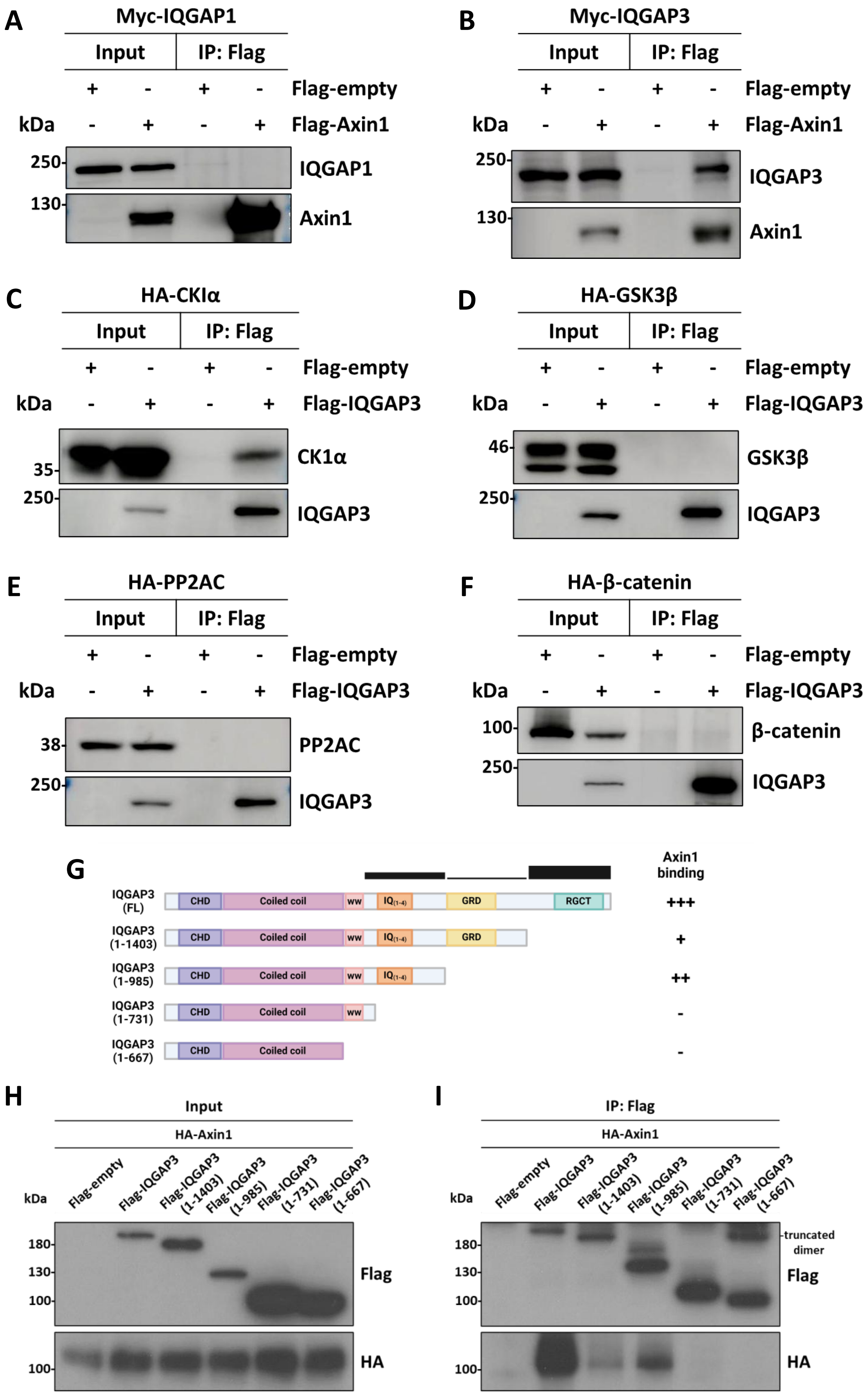

A
Myc-IQGAP1
Input
IP: Flag
Flag-empty
Flag-Axin1
kDa
250-
130-
IQGAP1
Axin1
B
Myc-IQGAP3
Input
IP: Flag
Flag-empty
Flag-Axin1
IQGAP3
Axin1
C
HA-CKIα
Flag-IQGAP3
35-
CK1α
IQGAP3
D
HA-GSK3β
46-
GSK3β
E
HA-PP2AC
38-
PP2AC
F
HA-β-catenin
100-
β-catenin
G
Axin1 binding
IQGAP3 (FL)
IQGAP3 (1-1403)
IQGAP3 (1-985)
IQGAP3 (1-731)
IQGAP3 (1-667)
CHD
Coiled coil
WW
IQ(1-4)
GRD
RGCT
+++
+
++
-
-
H
Input
HA-Axin1
Flag-empty
Flag-IQGAP3
Flag-IQGAP3 (1-1403)
Flag-IQGAP3 (1-985)
Flag-IQGAP3 (1-731)
Flag-IQGAP3 (1-667)
180-
130-
100-
Flag
HA
I
IP: Flag
truncated dimer

**Fig. S3: Exogenous immunoprecipitation.** (A) Coimmunoprecipitation of Myc-IQGAP1 with Flag-Axin1. (B) Coimmunoprecipitation of Myc-IQGAP3 with Flag-Axin1. (C) Coimmunoprecipitation of HA-CK1α with Flag-IQGAP3. (D) Coimmunoprecipitation of HA-GSK3β with Flag-IQGAP3. (E) Coimmunoprecipitation of HA-PP2AC with Flag-IQGAP3. (F) Coimmunoprecipitation of HA-β-catenin with Flag-IQGAP3.

**IQGAP3 binds to Axin1 c-terminal domain via its IQ and RGCT domains.** (G) Mapping of Axin1 binding site within IQGAP3. (H) immunoblot of input protein expressing Flag-IQGAP3 C-terminal truncations with HA-Axin1. (I) Immunoblot of Flag-Immunoprecipitated protein.

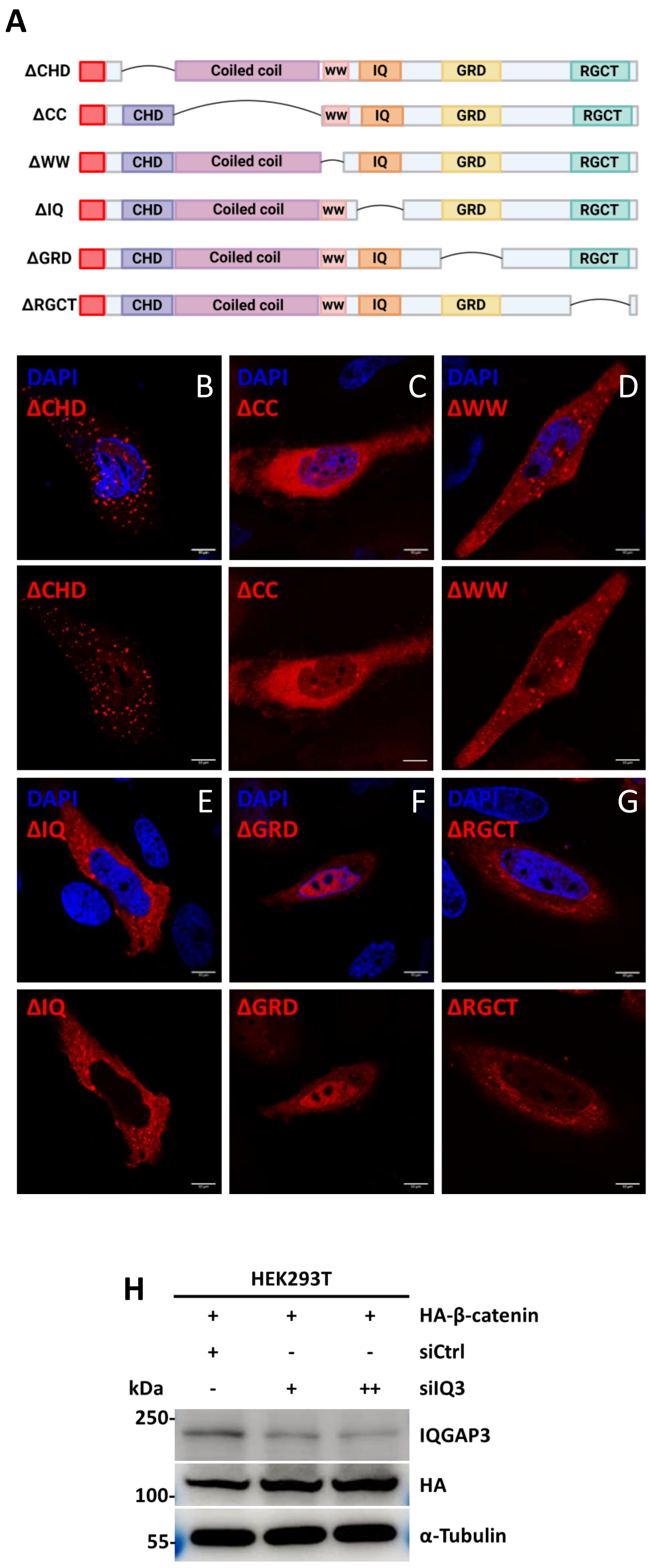
A
ΔCHD
ΔCC
ΔWW
ΔIQ
ΔGRD
ΔRGCT
CHD
Coiled coil
ww
IQ
GRD
RGCT
DAPI
B
C
D
E
F
G
ΔIQ
ΔGRD
H
HEK293T
HA-β-catenin
siCtrl
siIQ3
kDa
+ + +
+ - -
- + ++
250-
100-
55-
IQGAP3
HA
α-Tubulin

**Fig. S4: Microscopy images of IQGAP3 deletion constructs and Immunoblot of L cells and L-Wnt3a cell lysates, and MKN28 TurboID-IQGAP3 induction and biotinylation** (A) mCherry2-IQGAP3 deletion constructs. HeLa cells were transfected with the following constructs: (B) mCherry2-IQGAP3ΔCHD, (C) mCherry2-IQGAP3ΔCC, (D) mCherry2-IQGAP3ΔWW, (E) mCherry2-IQGAP3ΔIQ, (F) mCherry2-IQGAP3ΔGRD, (G) mCherry2-IQGAP3ΔRGCT. (H) Validation of IQGAP3 knockdown using siIQGAP3.

A

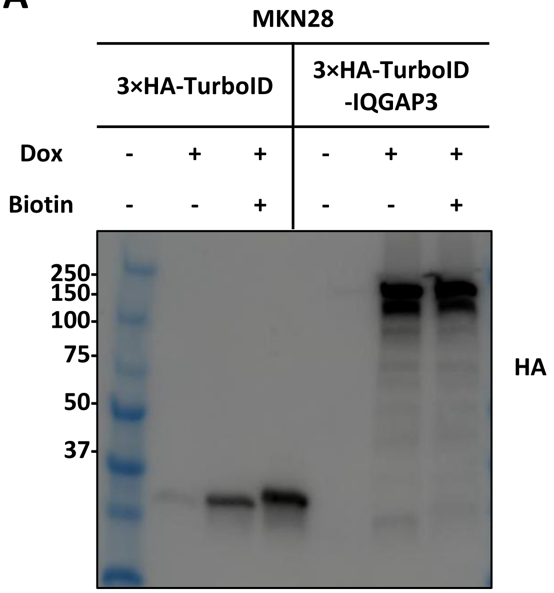


B

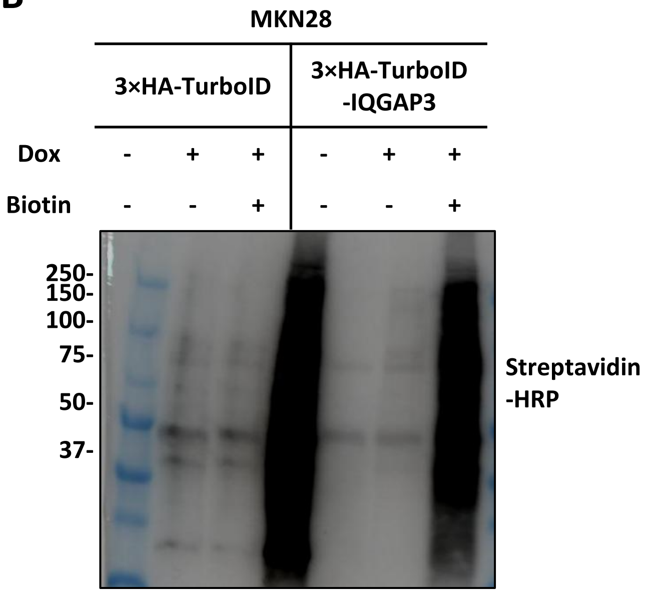


C

| | GS follow link to MSigDB | GS DETAILS | SIZE | ES | NES | NOM p-val | FDR q-val | FWER p-val | RANK AT MAX | LEADING EDGE |
|---|---|---|---|---|---|---|---|---|---|---|
| 1 | HALLMARK_G2M_CHECKPOINT | Details ... | 199 | 0.37 | 1.71 | 0.000 | 0.034 | 0.026 | 5107 | tags=56%, list=34%, signal=83% |
| 2 | HALLMARK_TGF_BETA_SIGNALING | Details ... | 53 | 0.45 | 1.65 | 0.008 | 0.031 | 0.048 | 2590 | tags=34%, list=17%, signal=41% |
| 3 | HALLMARK_NOTCH_SIGNALING | Details ... | 30 | 0.49 | 1.63 | 0.021 | 0.023 | 0.053 | 1248 | tags=33%, list=8%, signal=36% |
| 4 | HALLMARK_WNT_BETA_CATENIN_SIGNALING | Details ... | 38 | 0.47 | 1.60 | 0.015 | 0.027 | 0.080 | 2365 | tags=39%, list=16%, signal=47% |
| 5 | HALLMARK_HEDGEHOG_SIGNALING | Details ... | 30 | 0.45 | 1.46 | 0.053 | 0.075 | 0.256 | 2553 | tags=43%, list=17%, signal=52% |
| 6 | HALLMARK_MITOTIC_SPINDLE | Details ... | 198 | 0.30 | 1.41 | 0.014 | 0.097 | 0.381 | 4208 | tags=40%, list=28%, signal=55% |
| 7 | HALLMARK_ESTROGEN_RESPONSE_EARLY | Details ... | 173 | 0.29 | 1.31 | 0.022 | 0.171 | 0.620 | 3474 | tags=34%, list=23%, signal=43% |
| 8 | HALLMARK_E2F_TARGETS | Details ... | 200 | 0.28 | 1.31 | 0.030 | 0.159 | 0.643 | 4485 | tags=40%, list=30%, signal=56% |
| 9 | HALLMARK_ANGIOGENESIS | Details ... | 21 | 0.42 | 1.24 | 0.200 | 0.237 | 0.819 | 4268 | tags=62%, list=28%, signal=86% |
| 10 | HALLMARK_APICAL_JUNCTION | Details ... | 158 | 0.28 | 1.23 | 0.074 | 0.226 | 0.848 | 2240 | tags=19%, list=15%, signal=22% |
| 11 | HALLMARK_EPITHELIAL_MESENCHYMAL_TRANSITION | Details ... | 142 | 0.27 | 1.22 | 0.084 | 0.216 | 0.863 | 2692 | tags=32%, list=18%, signal=39% |
| 12 | HALLMARK_PI3K_AKT_MTOR_SIGNALING | Details ... | 89 | 0.29 | 1.22 | 0.133 | 0.202 | 0.869 | 5331 | tags=47%, list=35%, signal=73% |
| 13 | HALLMARK_INFLAMMATORY_RESPONSE | Details ... | 135 | 0.27 | 1.19 | 0.122 | 0.221 | 0.907 | 2294 | tags=24%, list=15%, signal=29% |
| 14 | HALLMARK_UV_RESPONSE_DN | Details ... | 124 | 0.24 | 1.07 | 0.285 | 0.457 | 0.991 | 2695 | tags=30%, list=18%, signal=36% |
| 15 | HALLMARK_ESTROGEN_RESPONSE_LATE | Details ... | 169 | 0.22 | 0.97 | 0.566 | 0.711 | 1.000 | 2410 | tags=21%, list=16%, signal=25% |
| 16 | HALLMARK_ANDROGEN_RESPONSE | Details ... | 87 | 0.24 | 0.97 | 0.517 | 0.679 | 1.000 | 2542 | tags=21%, list=17%, signal=25% |
| 17 | HALLMARK_PEROXISOME | Details ... | 85 | 0.22 | 0.89 | 0.669 | 0.884 | 1.000 | 1913 | tags=16%, list=13%, signal=19% |
| 18 | HALLMARK_HEME_METABOLISM | Details ... | 167 | 0.19 | 0.86 | 0.879 | 0.932 | 1.000 | 2080 | tags=17%, list=14%, signal=20% |
| 19 | HALLMARK_UV_RESPONSE_UP | Details ... | 142 | 0.19 | 0.83 | 0.855 | 0.926 | 1.000 | 1678 | tags=15%, list=11%, signal=16% |
| 20 | HALLMARK_CHOLESTEROL_HOMEOSTASIS | Details ... | 67 | 0.20 | 0.78 | 0.863 | 0.957 | 1.000 | 1955 | tags=18%, list=13%, signal=20% |
| 21 | HALLMARK_PROTEIN_SECRETION | | 92 | 0.15 | 0.61 | 1.000 | 0.993 | 1.000 | 4360 | tags=26%, list=29%, signal=36% |

D

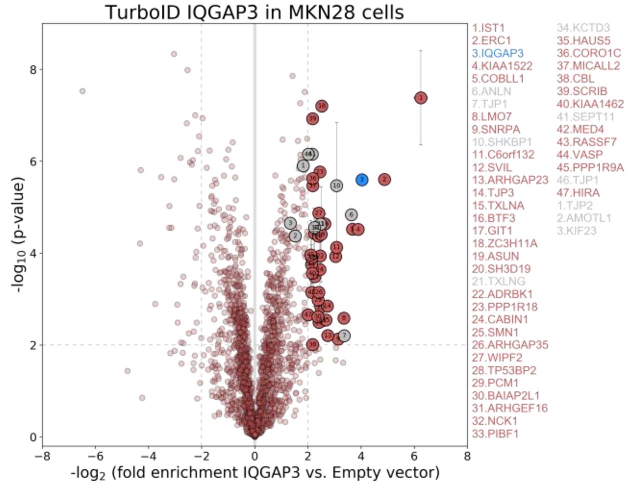


E

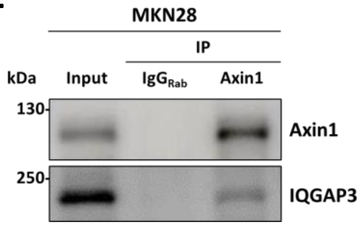


F

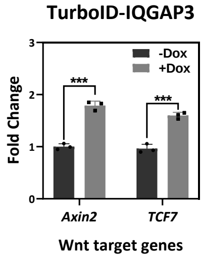

**Fig. S5: MKN28 TurboID-IQGAP3 expressing cells.** (A) Doxycycline induction of TurboID-IQGAP3 in MKN28 cells probed with anti-HA antibody (1:1000). (B) Biotin labelling of TurboID-IQGAP3 in MKN28 cells probed with streptavidin-HRP antibody (1:40,000). (C) Full GSEA ranking list from RNA-seq data of MKN28 cells expressing TurboID-IQGAP3. (D) of Volcano plot of biotinylated proteins found in MKN28 cells expressing TurboID-IQGAP3, (E) Validation of IQGAP3-Axin1 interaction in MKN28 lysate expressing TurboID-IQGAP3; Axin1 immunoprecipitation with MKN28 lysate and probing with Axin1 and IQGAP3 antibodies. (F) Validation of upregulated Wnt target genes *Axin2* and *TCF7.*

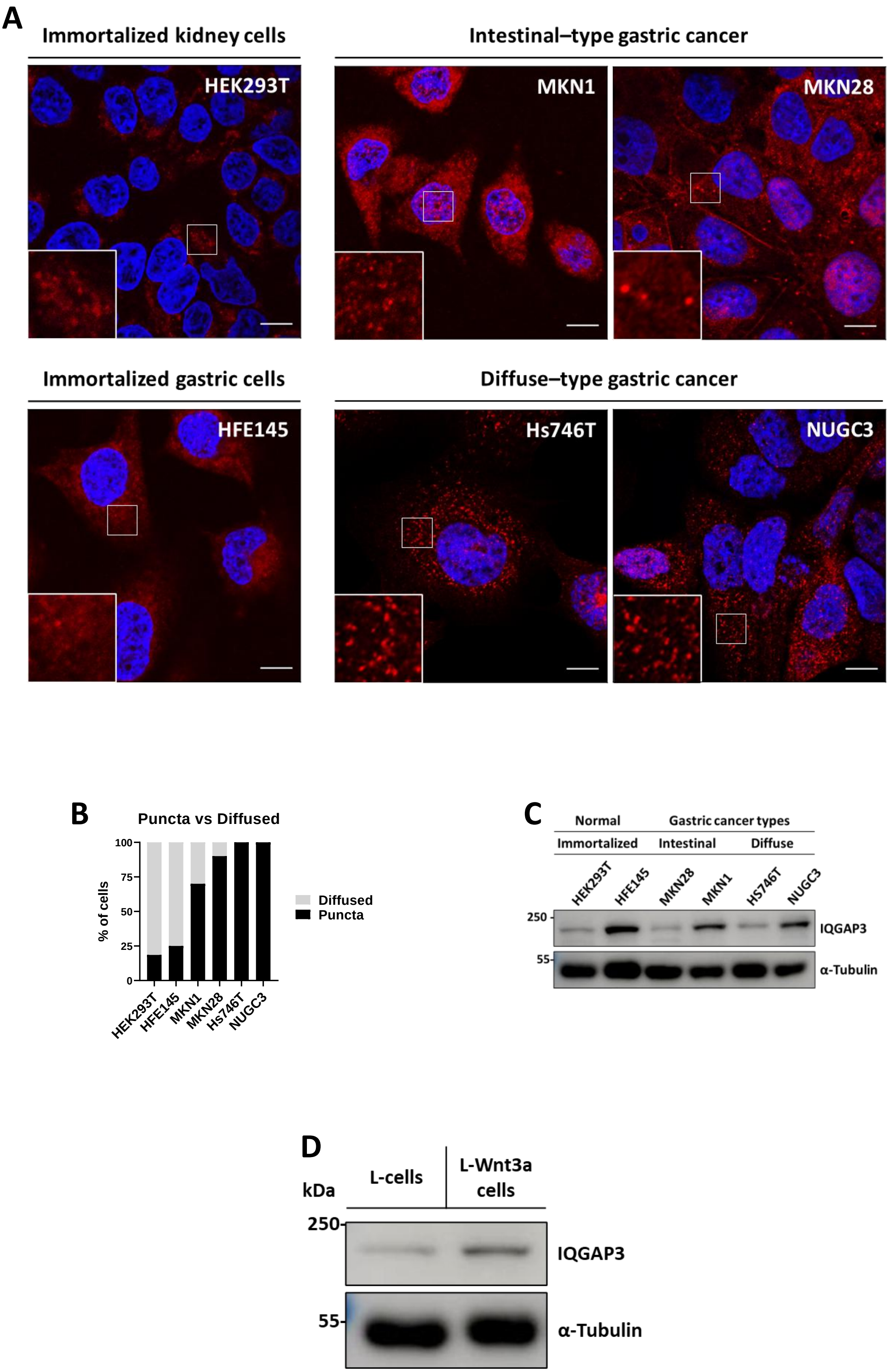
A
Immortalized kidney cells
HEK293T
Intestinal–type gastric cancer
MKN1
MKN28
Immortalized gastric cells
HFE145
Diffuse–type gastric cancer
Hs746T
NUGC3
B
Puncta vs Diffused
% of cells
100
75
50
25
0
HEK293T
HFE145
MKN1
MKN28
Hs746T
NUGC3
Diffused
Puncta
C
Normal
Gastric cancer types
Immortalized
Intestinal
Diffuse
HEK293T
HFE145
MKN28
MKN1
HS746T
NUGC3
250
55
IQGAP3
α-Tubulin
D
kDa
L-cells
L-Wnt3a cells
250
55
IQGAP3
α-Tubulin

**Fig. S6: MKN28 TurboID-IQGAP3 expressing cells.** (A) Immunofluorescence staining of HEK293T, HFE145, MKN1, MKN28, Hs746T and NUGC3 using anti-IQGAP3 (#PA5-56363) antibody, scale bars, 10µm. (B) Percentile of Puncta versus Diffused IQGAP3 morphology in HEK293T, HFE145, MKN1, MKN28, NUGC3 and HS746T cells from counting 30 cells. (C) Western blot analysis of HEK293T, HFE145, MKN1, MKN28, NUGC3 and HS746T cell lysates. (D) Immunoblot of L-cells and L-Wnt3a cells lysate probed for IQGAP3 and tubulin expressions.

**Fig. S7: Uncropped immunoblot of Figure 3A.** HeLa Tet-On lysates induced with Doxycycline. (A) Membrane probed with GFP (D5.1) Rabbit mAb #2956 (1:1000). (B) Membrane blot probed with IQGAP3 #25930-1-AP, (1:1000). (C) Membrane blot probed with α-tubulin #11224-1-AP, (1:10,000). First lane, Protein ladder: PageRuler™ Plus Prestained 10 to 250 kDa #26619. Resolved on 7.5% SDS-PAGE. Cropped image as shown in main manuscript are shown within the red box.

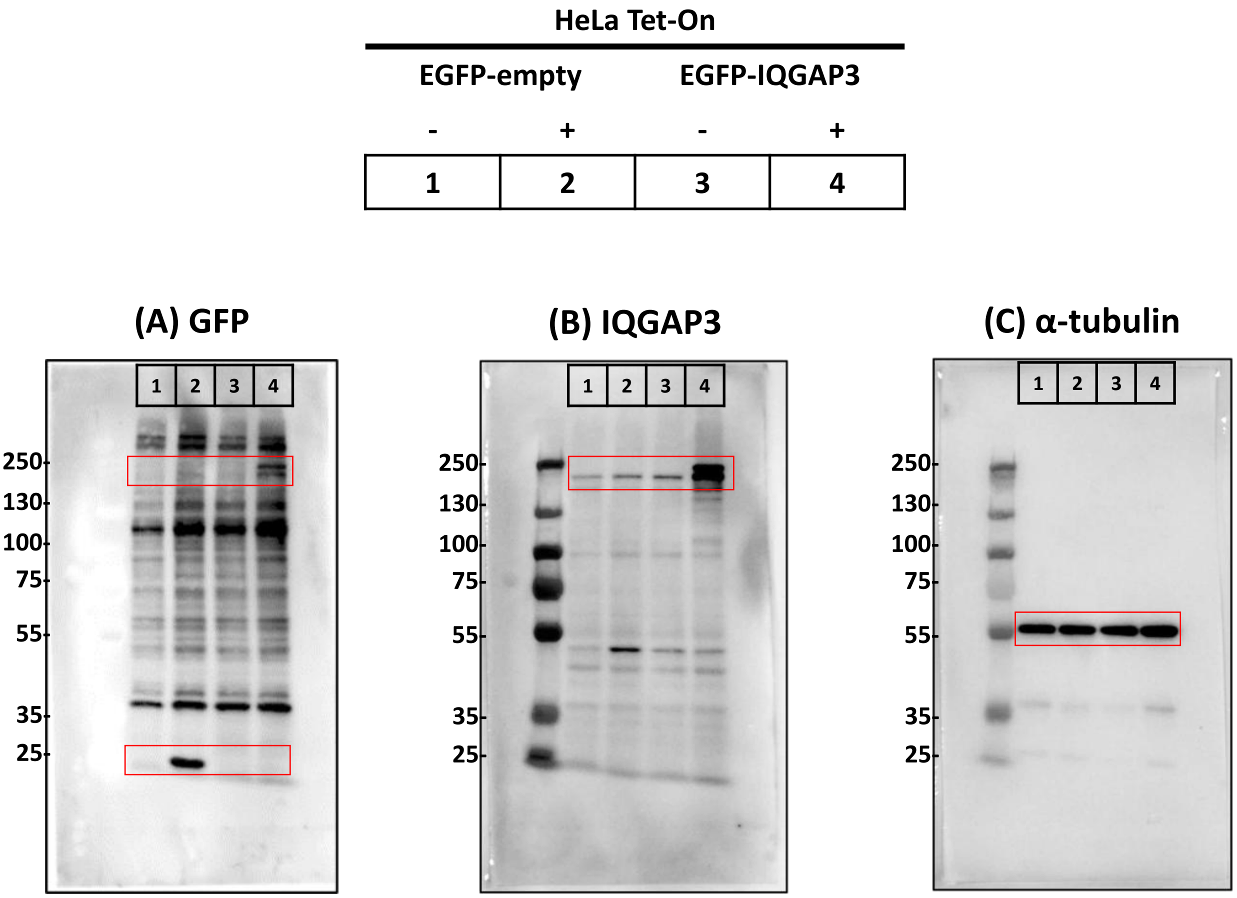

**Fig. S8: Uncropped immunoblot of Figure 5A.** IQGAP3 immunoprecipitation from HEK293 cell lysate using rabbit polyclonal anti-IQGAP3 #89-301-822, Antibody pair for IP (1:100). (A) Membrane blot probed with mouse polyclonal anti-IQGAP3 (#89-301-822, Antibody pair for WB) (1:1000). (B) Membrane blot probed with Axin1 #1C4E8 (1:1000). (C) Membrane blot probed with β-catenin antibody (D10A8) #8480, (1:1000). (D) Membrane blot probed with CK1α antibody (H-7) #sc-74582, (1:250) (E) Membrane blot probed with PP2Aα antibody (D10A8) #8480, (1:1000). (F) Membrane blot probed with GSK-3β antibody (3D10) #9832, (1:1000). (G) Membrane blot probed with PCM1 antibody (G2000) #5213, (1:1000). First and Fifth lane, Protein ladder: PageRuler™ Plus Prestained 10 to 250 kDa #26619. Resolved on 7.5% SDS-PAGE. Cropped image as shown in main manuscript are shown within the red box.

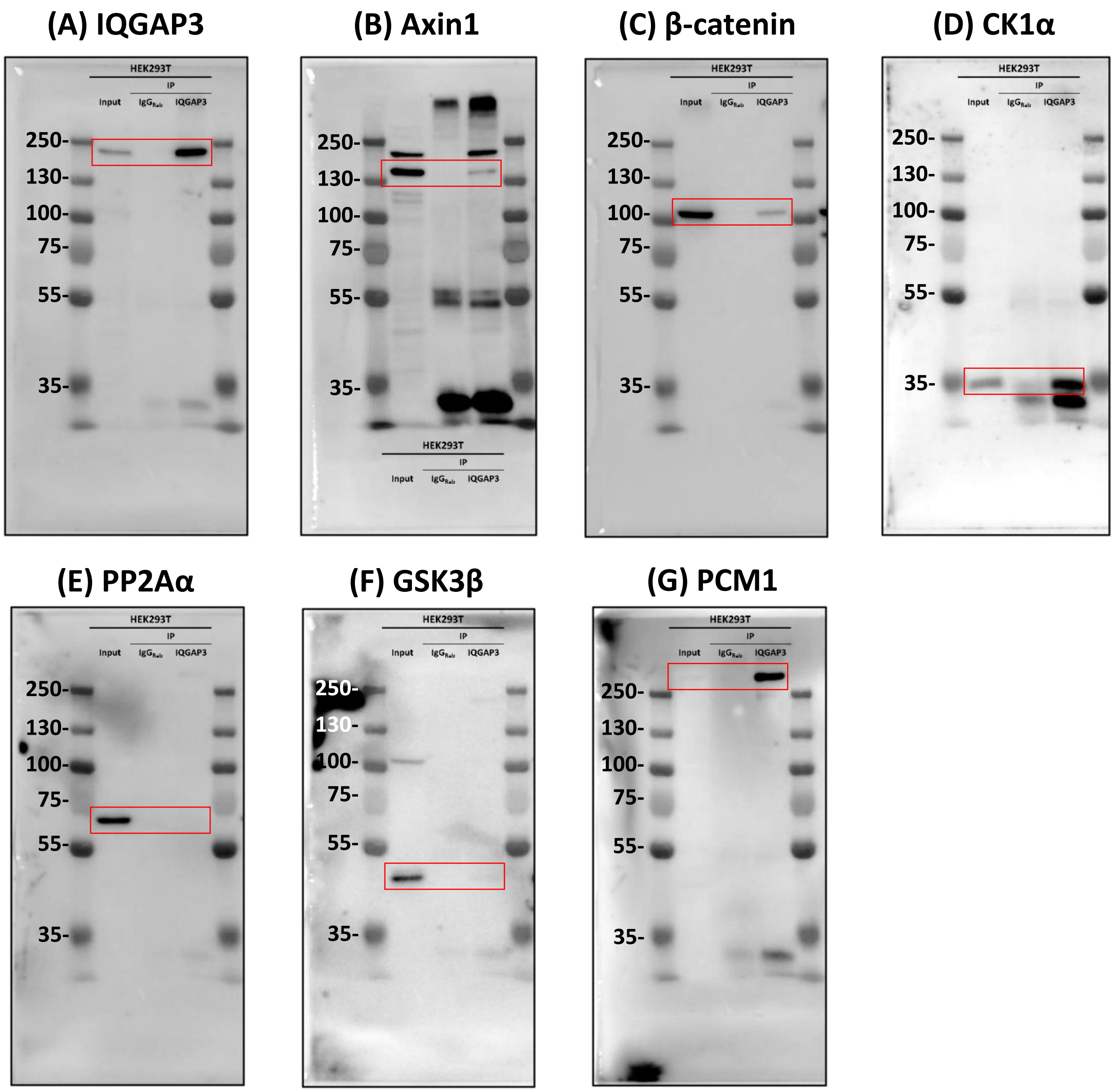

**Fig. S9: Uncropped immunoblot of Figure 5B.** Axin1 immunoprecipitated from HEK293T cell lysates using Axin1 Rabbit mAb (C76H11) #2087 (1:100). (A) Membrane blot probed with β-catenin antibody (D10A8) #8480, (B) Membrane blot probed with Axin1 (C76H11) #2087, (1:1000). (C) Membrane blot probed with IQGAP3 #25930-1-AP, (1:1000). (D) Membrane blot probed with GSKβ (D5C5Z) #12456, (1:1000). First lane, Protein ladder: PageRuler™ Plus Prestained 10 to 250 kDa #26619. Resolved on 7.5% SDS-PAGE. Cropped image as shown in main manuscript are shown within the red box.

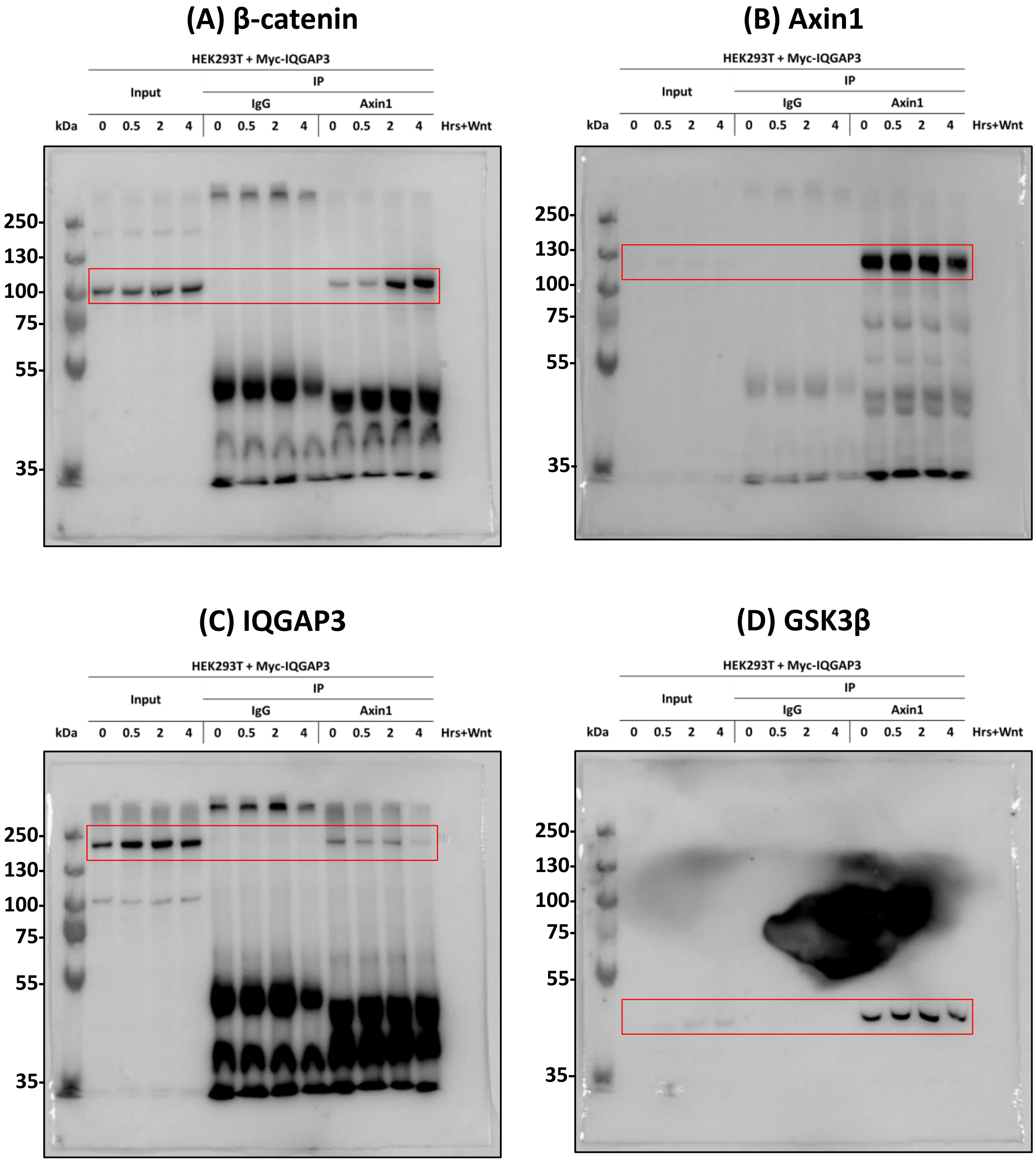

**Fig. S10: Uncropped immunoblot of Figure 5C.** IQGAP3 immunoprecipitated from HEK293T cell lysates using Myc-Tag (9B11) Mouse mAb #2276 (1:100). (A) Membrane blot probed with β-catenin antibody (D10A8) #8480, (1:1000). (B) Membrane blot probed with Axin1 (C76H11) #2087, (1:1000). (C) Membrane blot probed with IQGAP3 #25930-1-AP, (1:1000). (D) Membrane blot probed with IQGAP1 (D6E3J) #29016, (1:1000). First lane, Protein ladder: PageRuler™ Plus Prestained 10 to 250 kDa #26619. Resolved on 7.5% SDS-PAGE. Cropped image as shown in main manuscript are shown within the red box.

**(A) β-catenin**

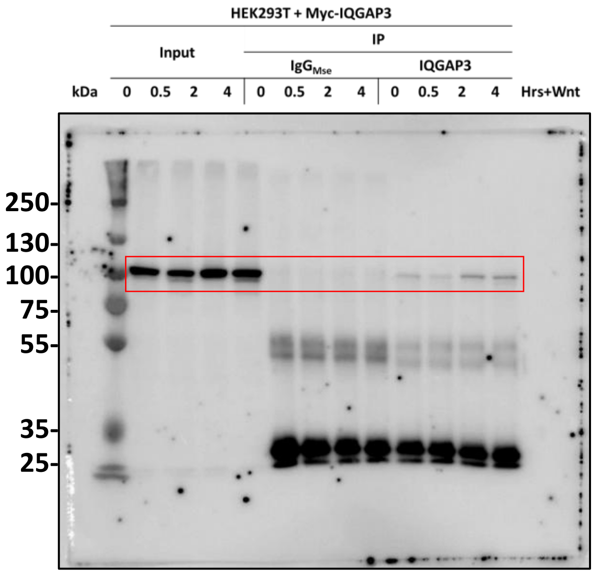


**(B) Axin1**

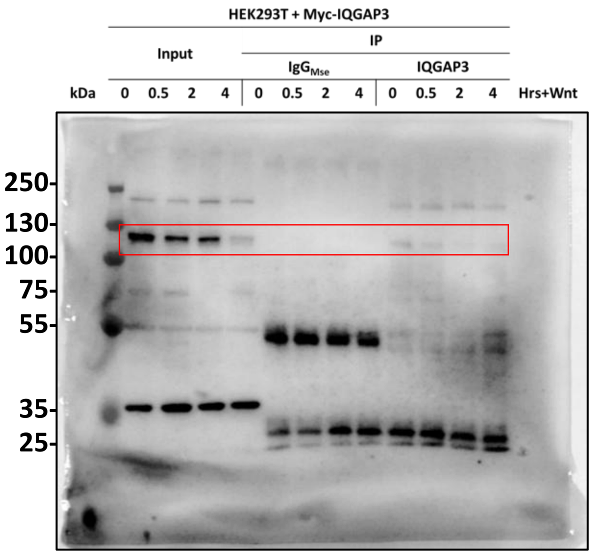


**(C) IQGAP3**

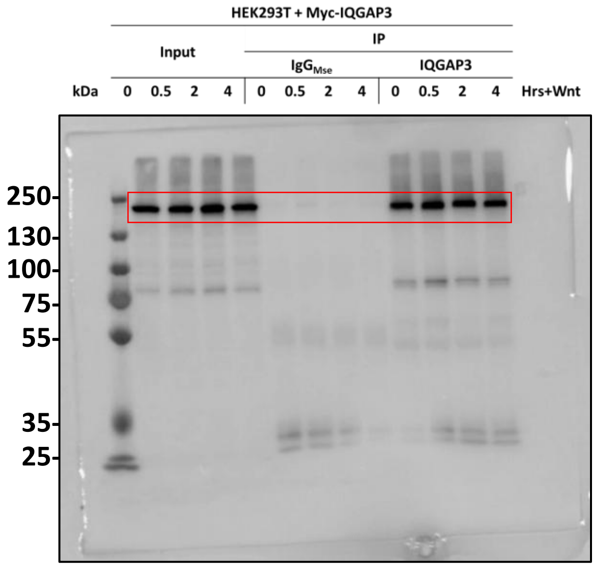


**(D) IQGAP1**

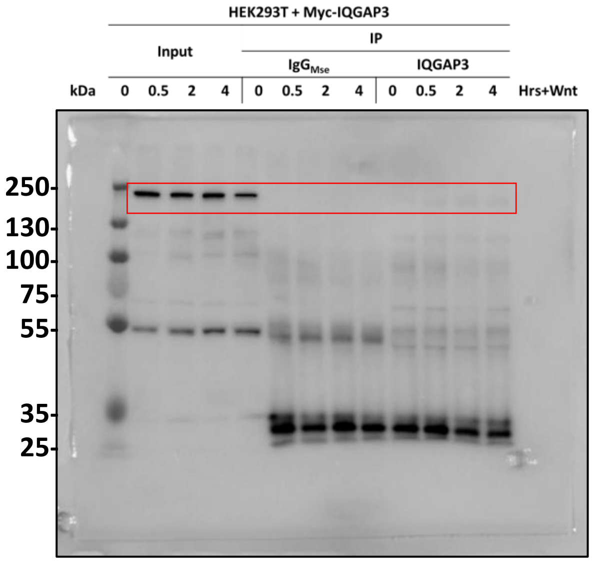

**Fig. S11: Uncropped immunoblot of Figure 5D.** Flag-IQGAP3 immunoprecipitated with Flag M2 Mouse mAb Antibody, #F3165, (1:100). (A and C) Membrane blot probed with HA-Tag (C29F4) Rabbit mAb #3724 antibody (1:1000). (B and D) Membrane blot probed with anti-FLAG rabbit antibody #F7425 (1:1000). First lane, Protein ladder: PageRuler™ Plus Prestained 10 to 250 kDa #26619. Resolved on 7.5% SDS-PAGE. Cropped image as shown in main manuscript are shown within the red box.

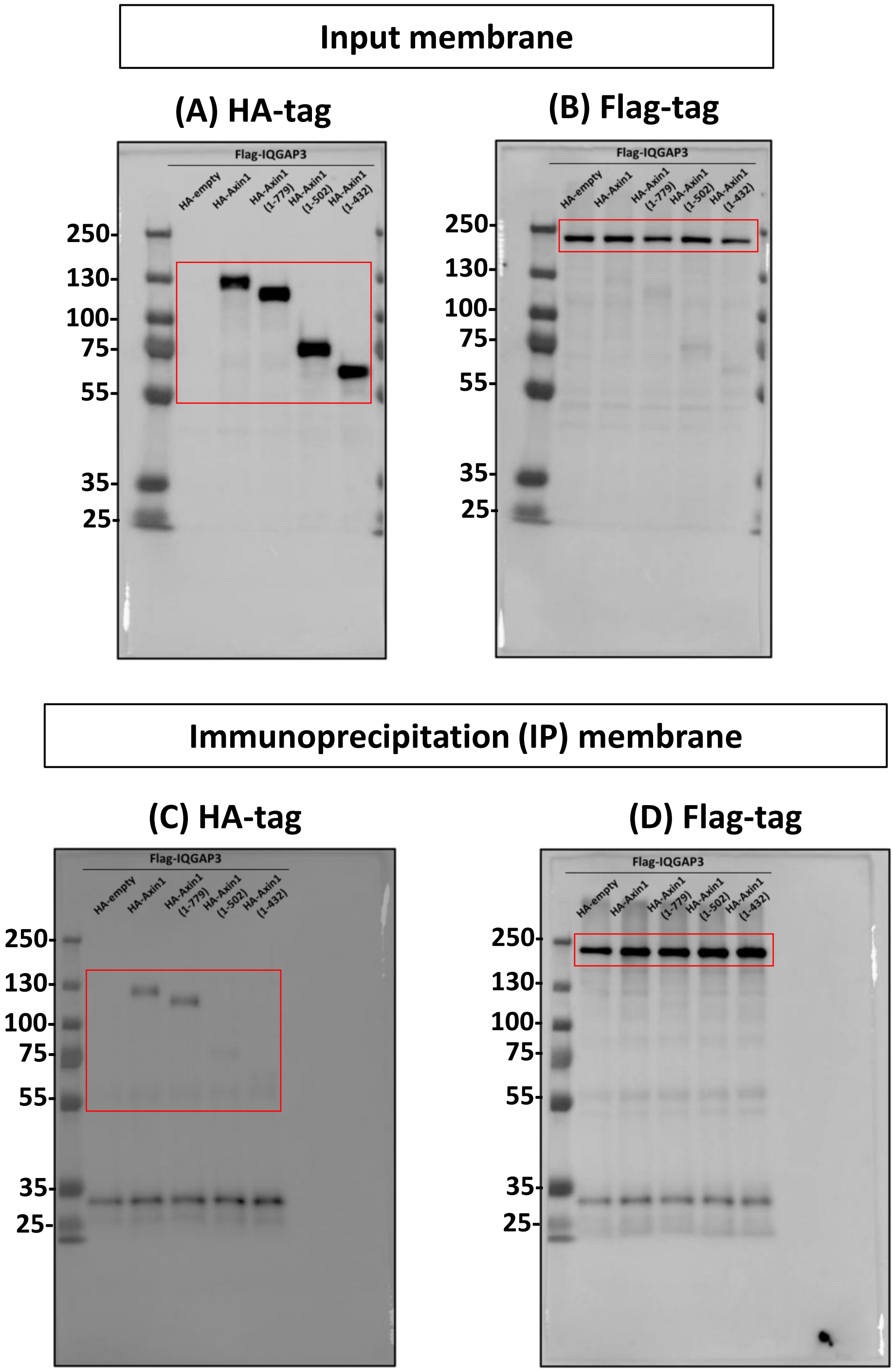

**Fig. S12: Uncropped immunoblot of Figure 5E.** Flag-Axin1 immunoprecipitated with Flag M2 Flag M2 Mouse mAb Antibody, #F3165, (1:100). (A and D) Membrane blot probed with anti-FLAG rabbit antibody #F7425 (1:1000). (B and E) Membrane blot probed with IQGAP3 #25930-1-AP, (1:1000). (C and F) Membrane blot probed with HA-Tag (C29F4) Rabbit mAb #3724 antibody (1:1000). First lane, Protein ladder: PageRuler™ Plus Prestained 10 to 250 kDa #26619. Resolved on 7.5% SDS-PAGE. Cropped image as shown in main manuscript are shown within the red box.

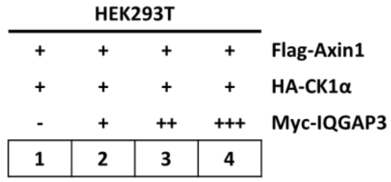


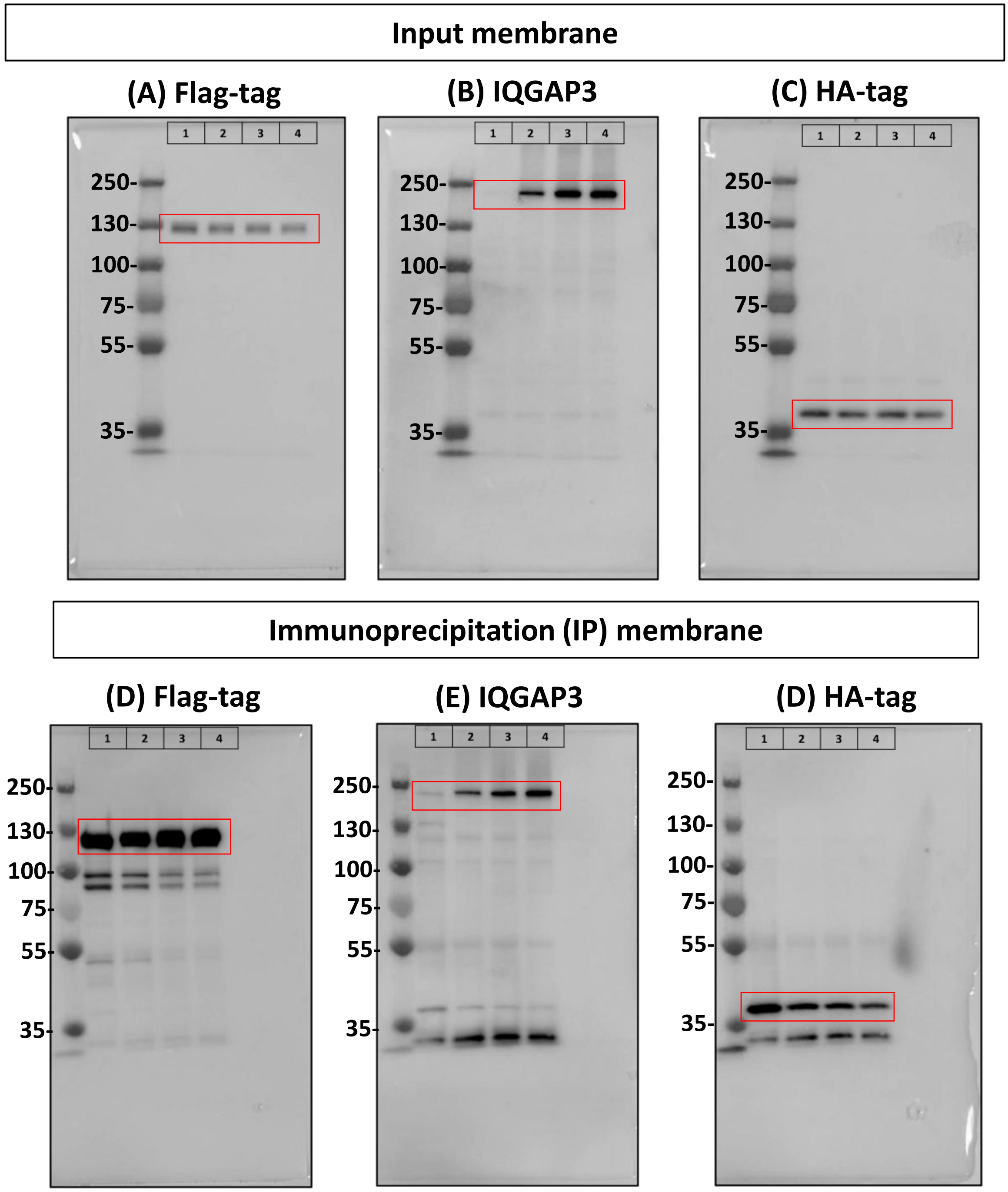

**Fig. S13: Uncropped immunoblot of Figure 5F.** Axin1 immunoprecipitated with Axin1 (C76H11) Rabbit mAb #2087 (1:50). (A and D) Membrane blot probed with CK1α Mouse Antibody (H-7) #sc-74582 (1:1000). (B and E) Membrane blot probed with Axin1 (C76H11) Rabbit mAb #2087 (1:1000). (C and F) Membrane blot with Myc-Tag (71D10) Rabbit mAb #2278 (1:1000). First lane, Protein ladder: PageRuler™ Plus Prestained 10 to 250 kDa #26619). Resolved on 10% SDS-PAGE. Cropped image as shown in main manuscript are shown within the red box.

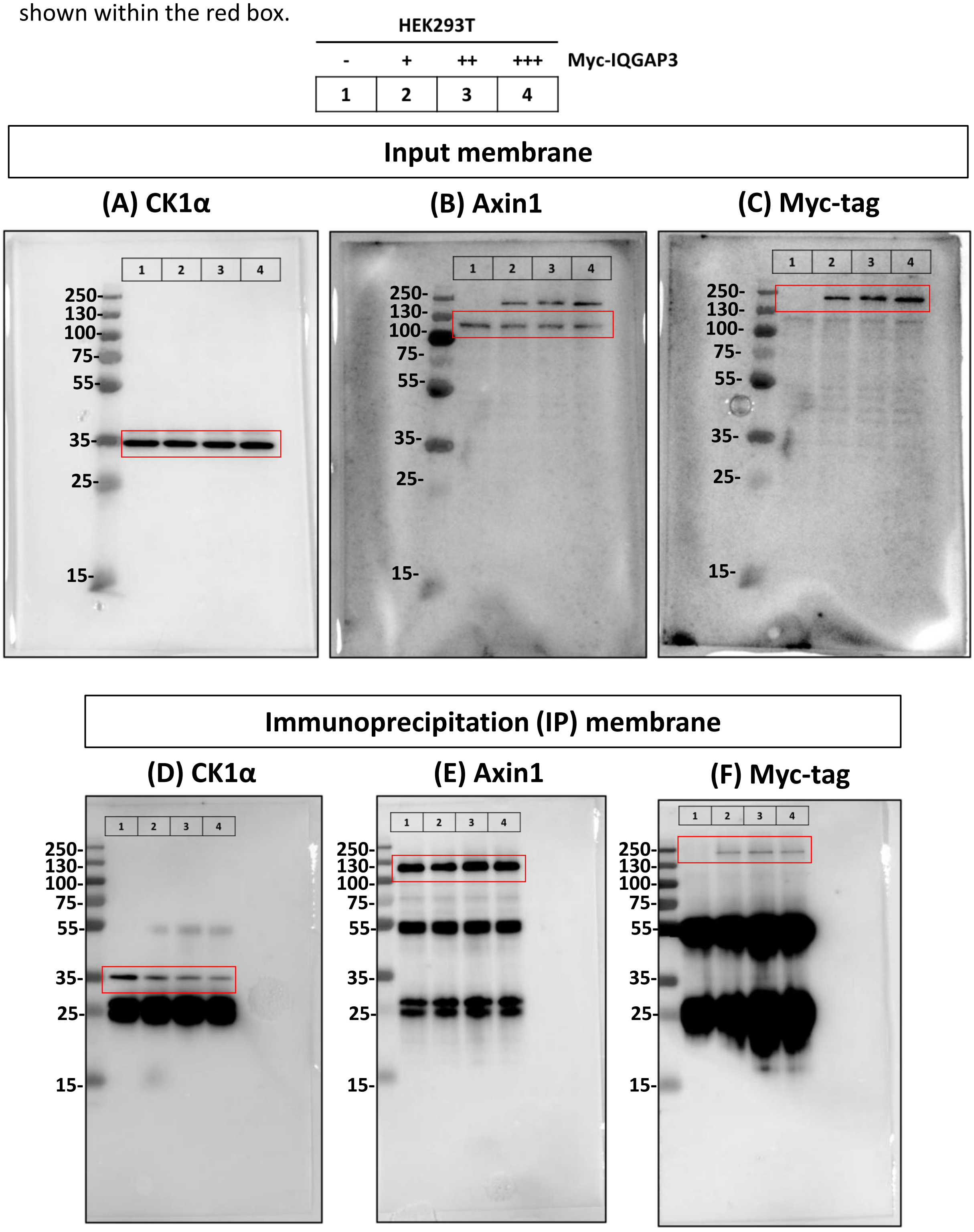

**Fig. S14: Uncropped Immunoblot of Figure 6B.** (A) Membrane blot probed with Non-phospho (Active) β-Catenin (Ser45) (D2U8Y) #19807, (1:1000) (B) Membrane blot probed with HA-Tag (C29F4) #3724, (1:1000). (C) Membrane blot probed with IQGAP3 #25930-1-AP, (1:1000). (D) Membrane blot probed with α-tubulin #11224-1-AP, (1:10,000). Prestained Protein Ladder - Broad molecular weight (10 - 245 kDa). Resolved on 7.5% SDS-PAGE. Cropped image as shown in main manuscript are shown within the red box.

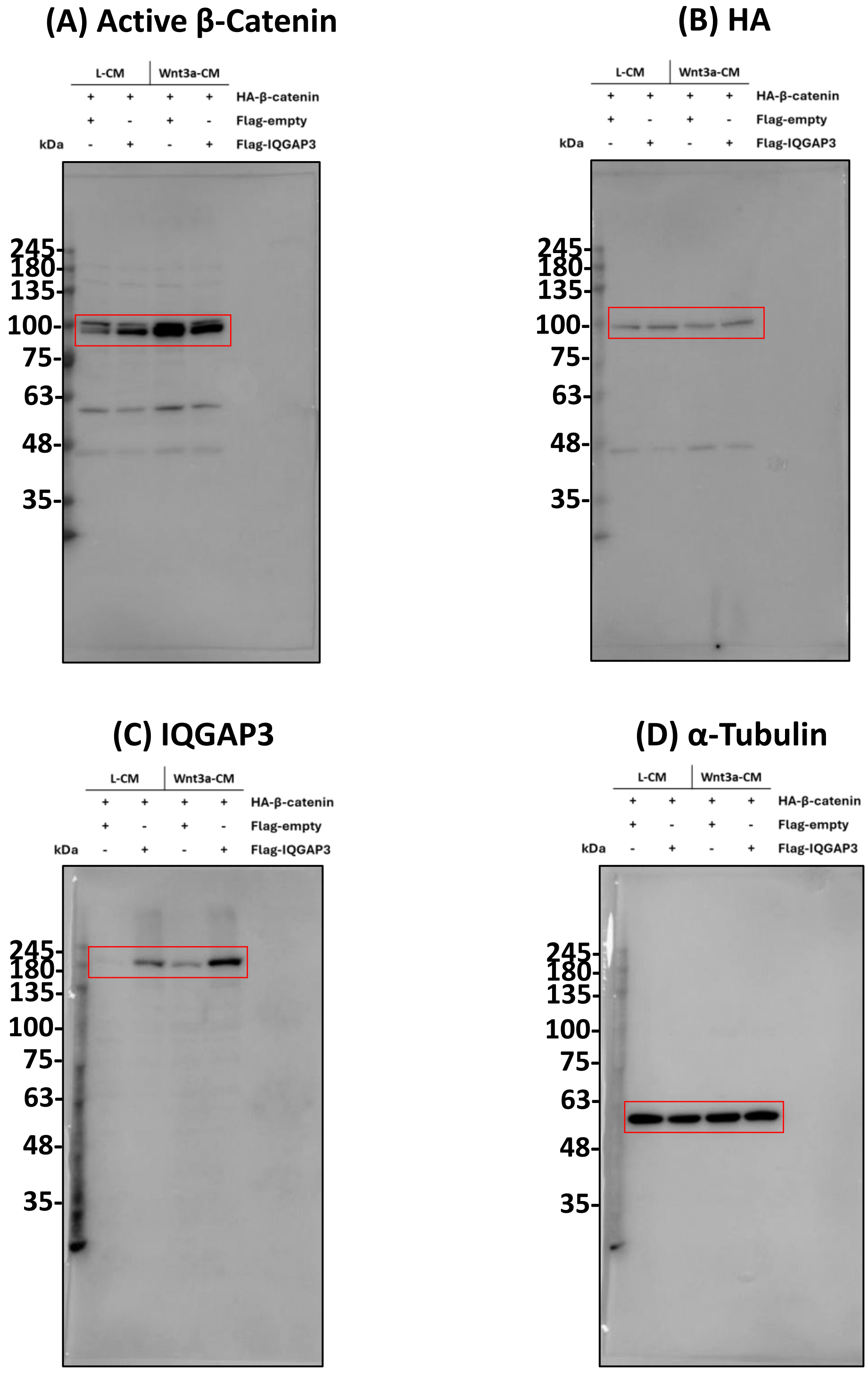

**Fig. S15: Uncropped immunoblot of Figure 6D and 7A** (A) Membrane blot probed with mCherry antibody #26765-1-AP, (1:1000). (B) Membrane blot probed with α-tubulin #11224-1-AP, (1:10,000). First lanes, Protein ladder PageRuler™ Plus Prestained 10 to 250 kDa #26619. Resolved on 7.5% SDS-PAGE. Cropped image as shown in main manuscript are shown within the red box. (C) Membrane blot probed with IQGAP3 #25930-1-AP, (1:1000). (D) Membrane blot probed with α-tubulin #11224-1-AP, (1:10,000). Prestained Protein Ladder - Broad molecular weight (10 - 245 kDa). Resolved on 7.5% SDS-PAGE. Cropped image as shown in main manuscript are shown within the red box.

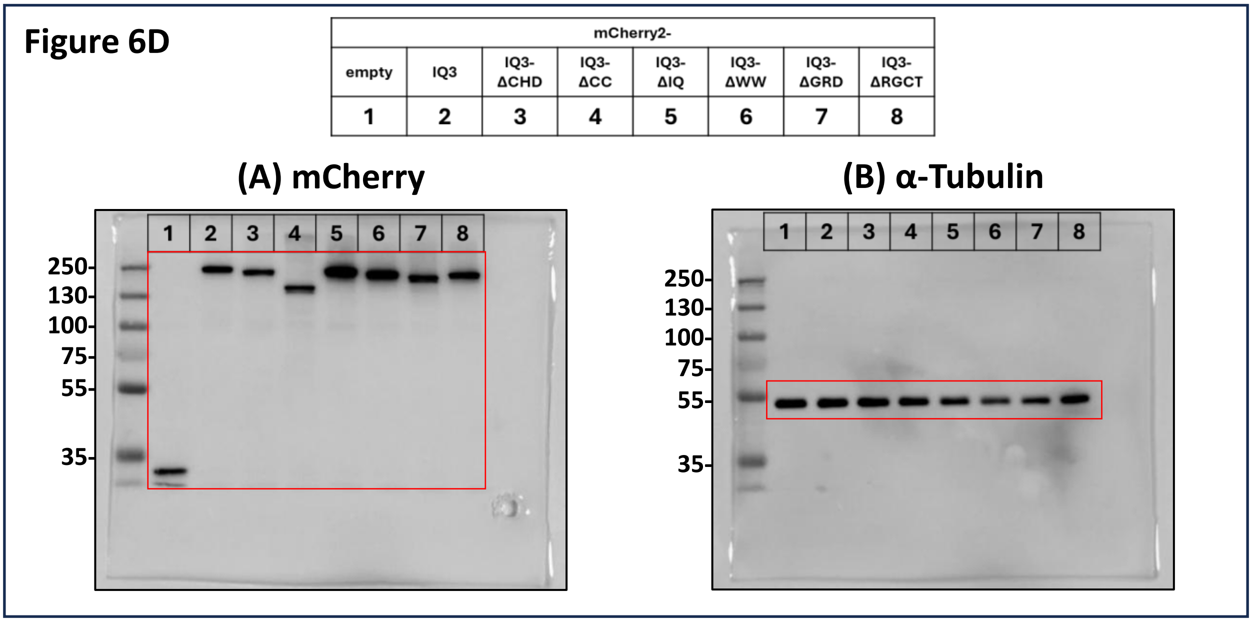


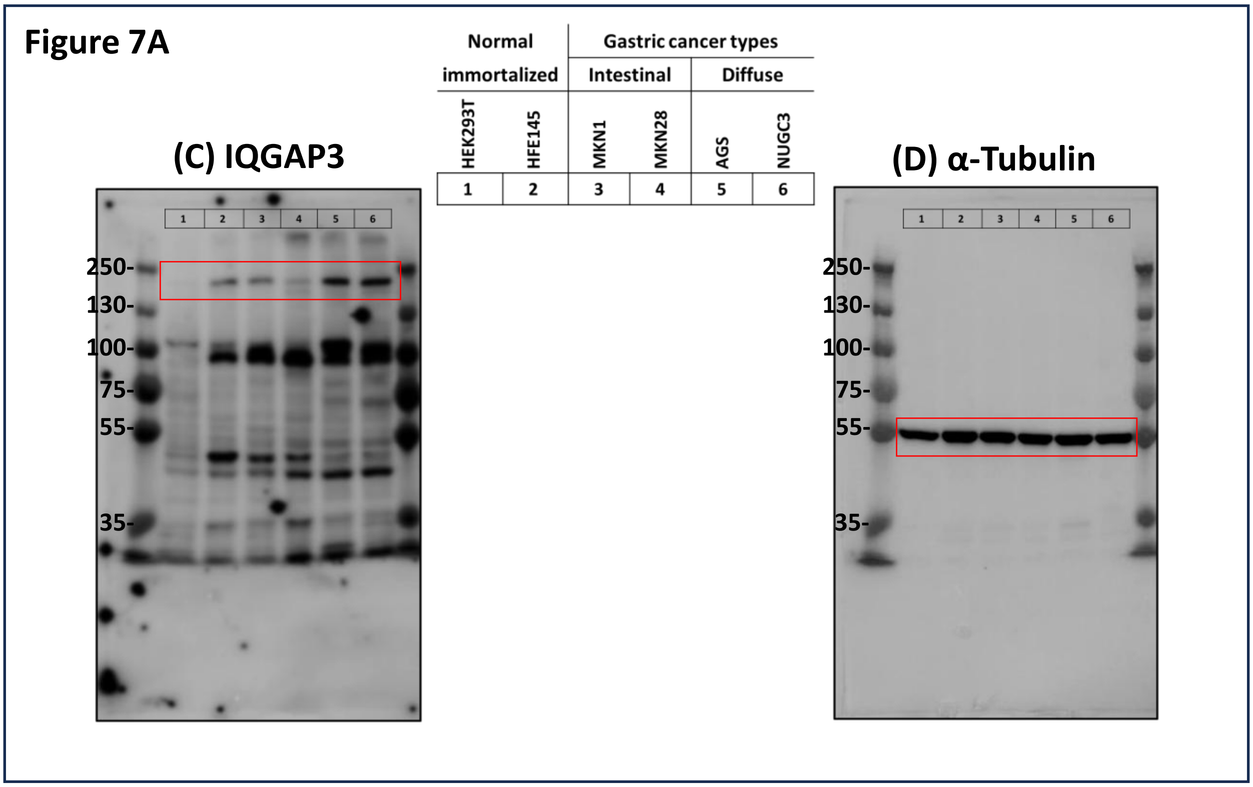

**Fig. S16: Uncropped immunoblot of Figure 7B and Figure 8A.** (A and D) Membrane blot probed with β-catenin antibody (D10A8) #8480, (1:1000). (B and E) Membrane blot probed with IQGAP3 #25930-1-AP, (1:1000). (C) Membrane blot probed with GAPDH (14C10) Rabbit mAb #2118 (1:1000). (F) Membrane blot probed with β-actin #, (1:10,000). First lanes, Protein ladder PageRuler™ Plus Prestained 10 to 250 kDa #26619. Resolved on 7.5% SDS-PAGE. Cropped image as shown in main manuscript are shown within the red box.

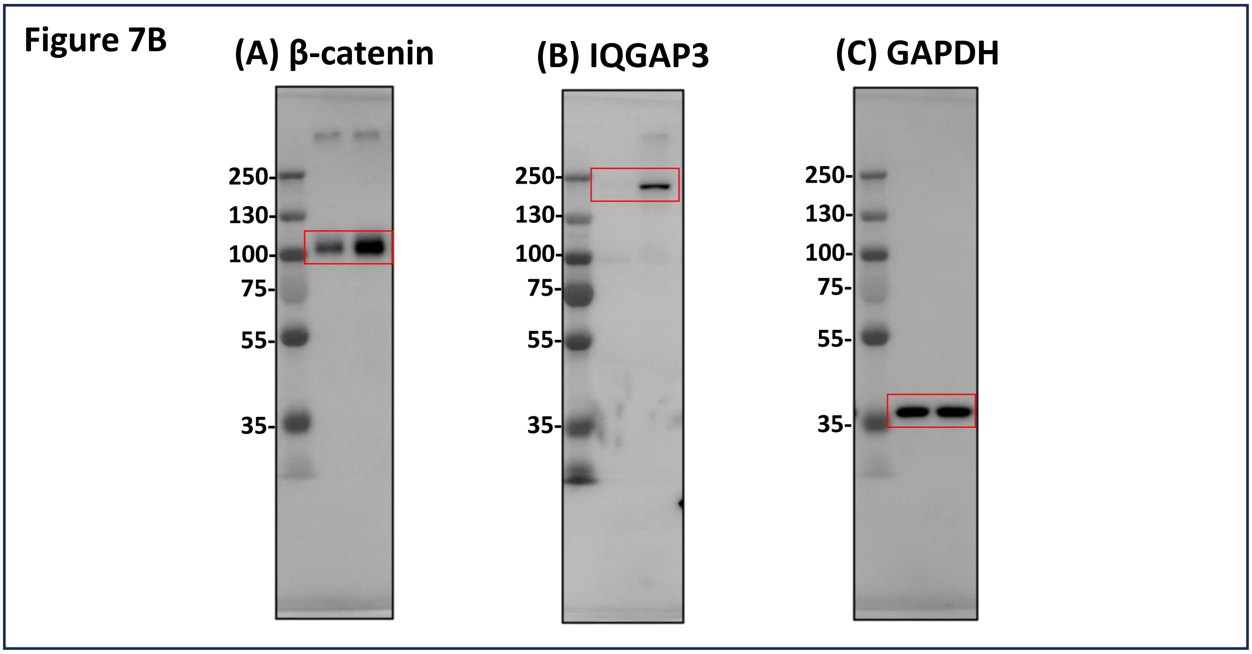


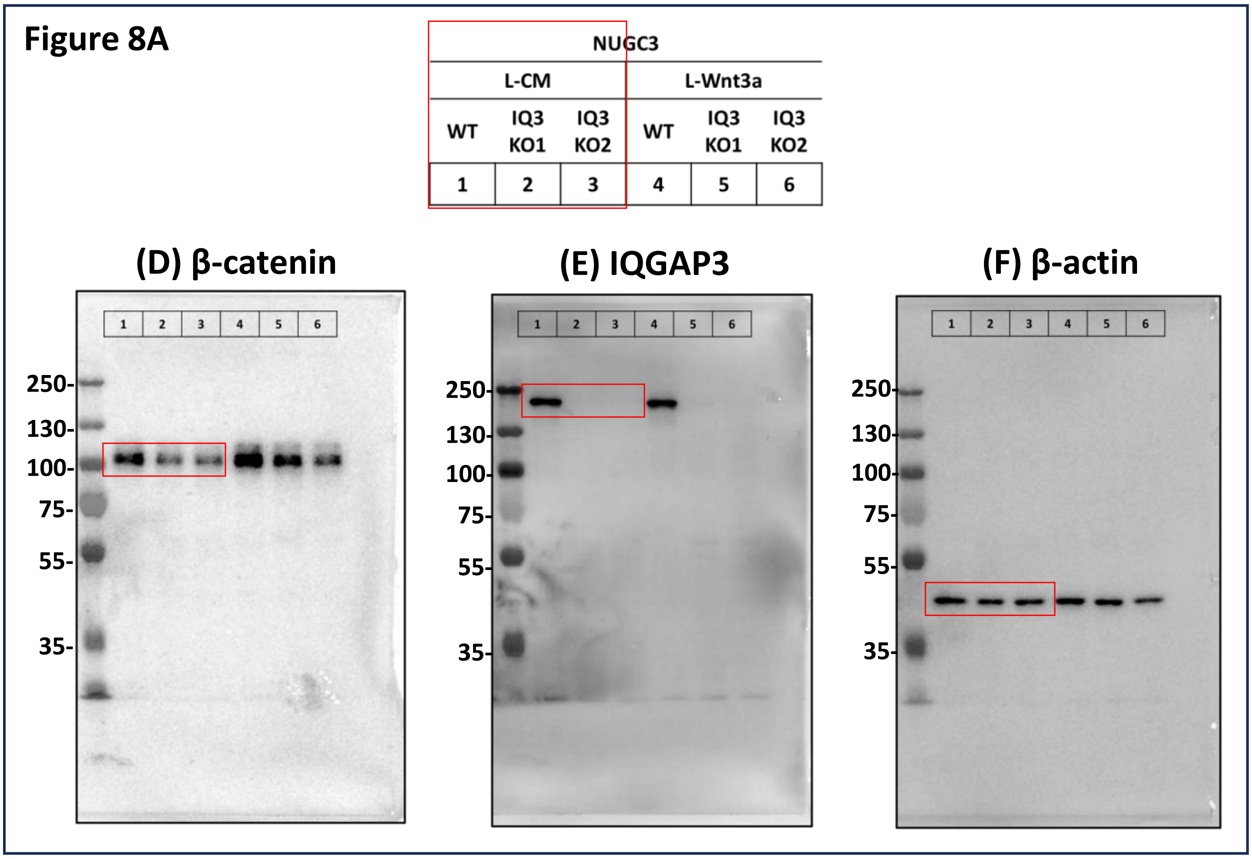

**Fig. S17: Uncropped immunoblot of Figure 8F.** (A) Membrane blot probed with β-catenin antibody (D10A8) #8480, (1:1000). (B) Membrane blot probed with IQGAP3 #25930-1-AP, (1:1000). (C) Membrane blot probed with α-tubulin #11224-1-AP, (1:10,000). Resolved on 7.5% SDS-PAGE. Cropped image as shown in main manuscript are shown within the red box.

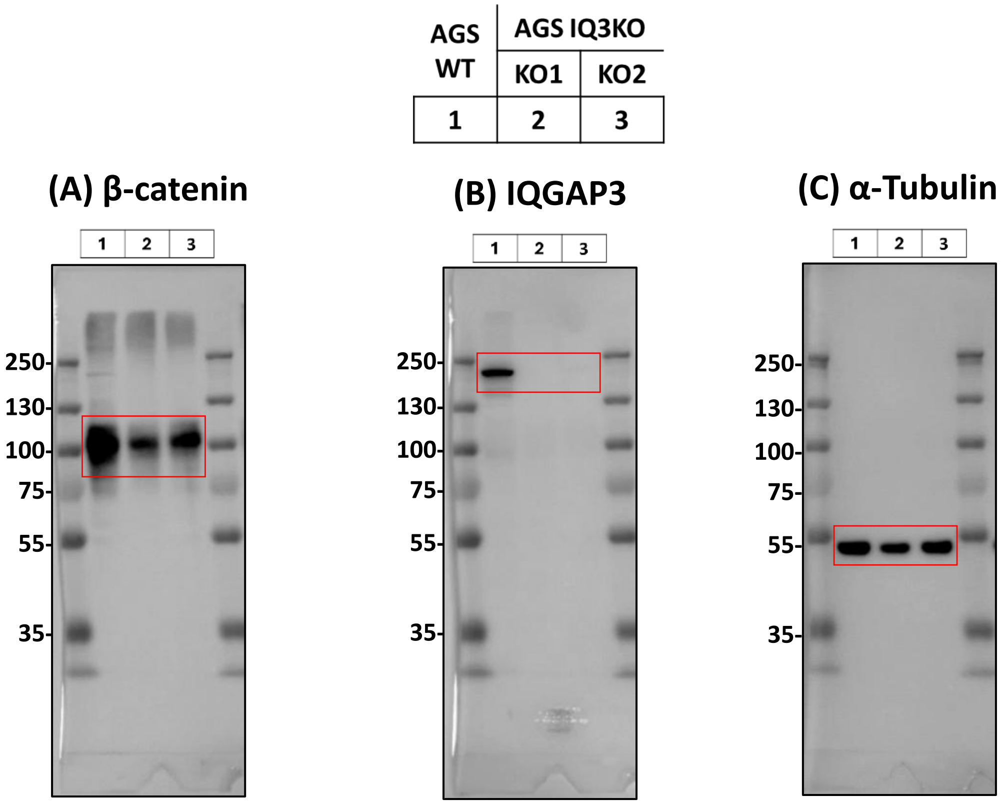

**Fig. S18: Uncropped immunoblot of Figure 9B.** (A) Membrane blot probed with β-catenin antibody (D10A8) #8480, (1:1000). (B) Membrane blot probed with Non-phospho (Active) β-Catenin (Ser45) (D2U8Y) #19807, (1:1000). (C) Membrane blot probed with Axin1 (C76H11) #2087, (1:1000). (D) Membrane blot probed with IQGAP3 #25930-1-AP, (1:1000). (E) Membrane blot probed with β-actin #4967 (1:10,000). First lane, Protein ladder PageRuler™ Plus Prestained 10 to 250 kDa #26619. Resolved on 7.5% SDS-PAGE. Cropped image as shown in main manuscript are shown within the red box.

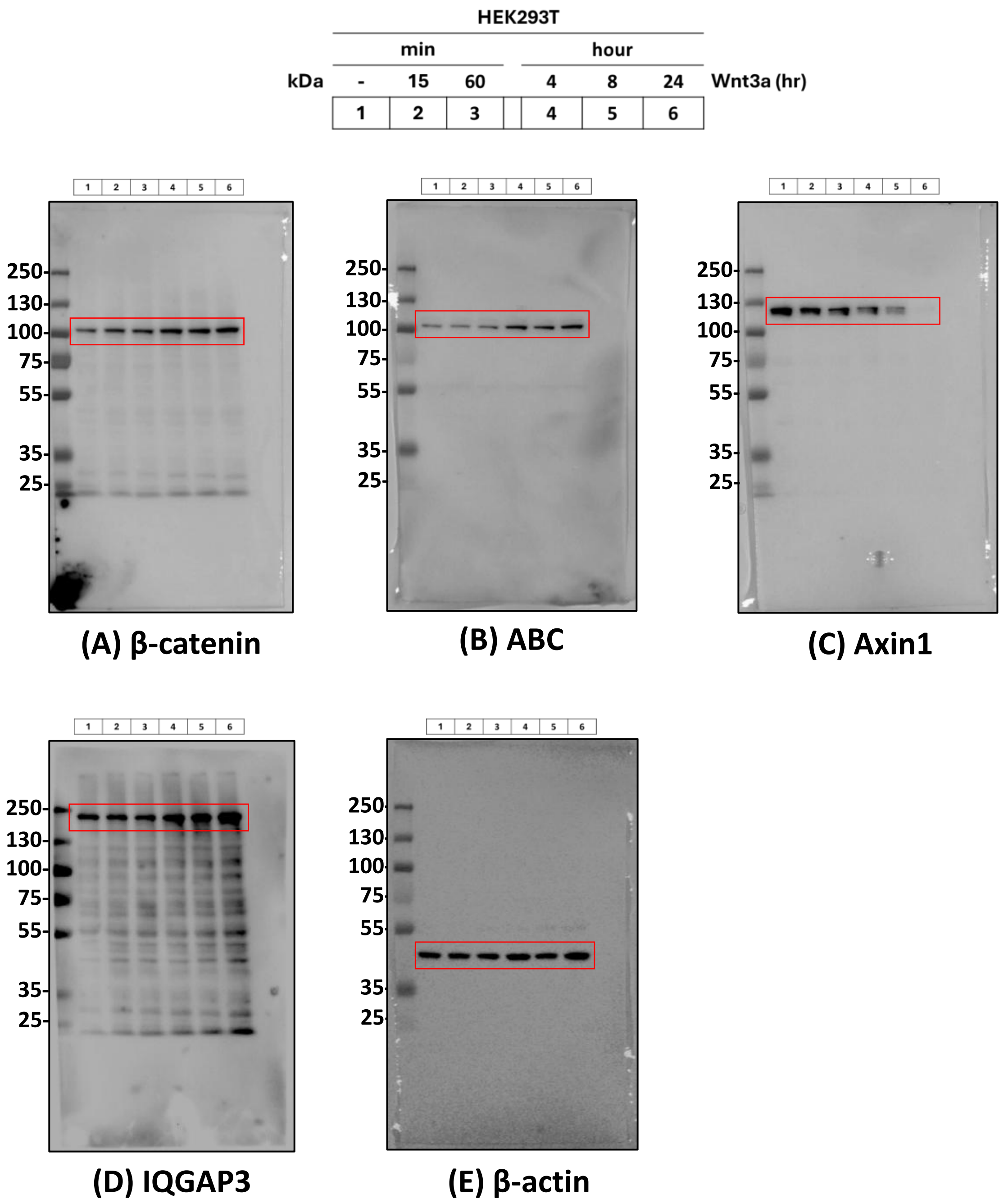

**Fig. S19: Uncropped immunoblot of Figure 9F.** (A) Membrane blot probed with β-catenin antibody (D10A8) #8480, (1:1000). (B) Membrane blot probed with IQGAP3 #25930-1-AP, (1:1000). (C) Membrane blot probed with Axin1 (C76H11) #2087, (1:1000). (D) Membrane blot probed with α-tubulin #11224-1-AP, (1:10,000). First lane, Protein ladder PageRuler™ Plus Prestained 10 to 250 kDa #26619. Resolved on 7.5% SDS-PAGE.

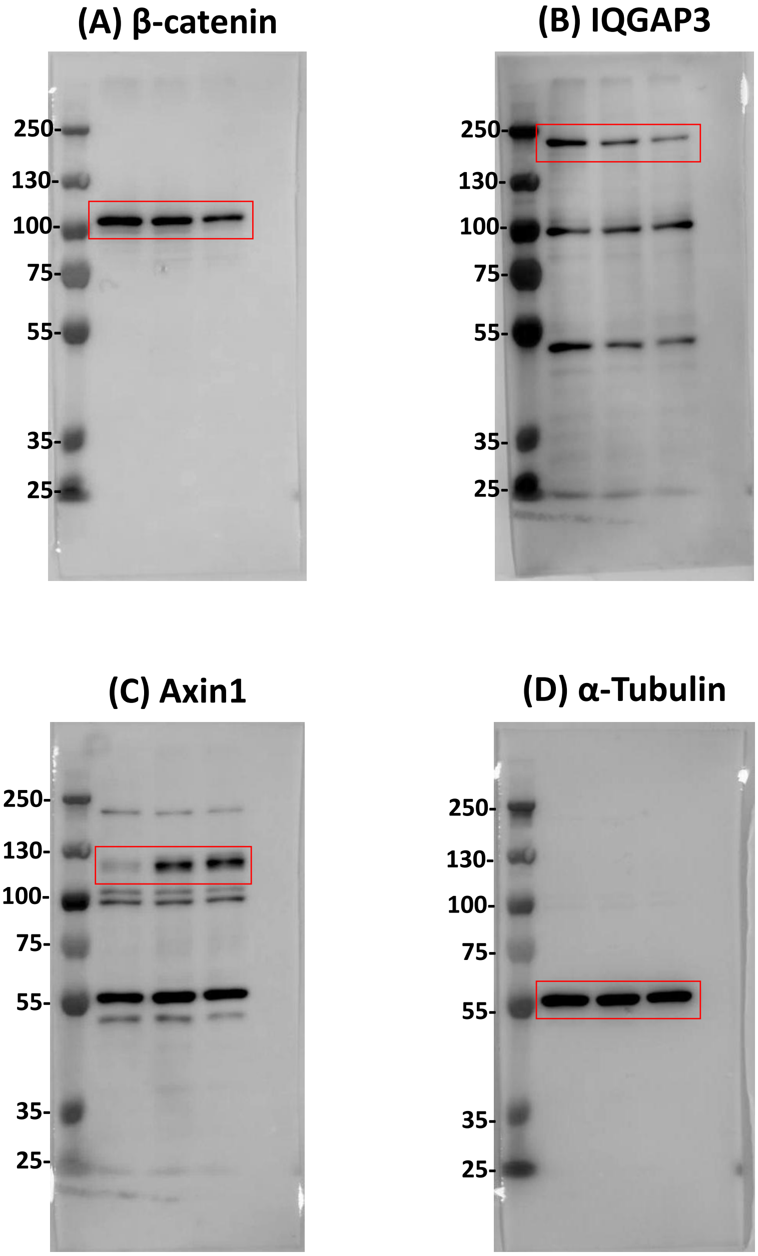
